# A continuous-time microparasite model incorporating infection intensity and parasite aggregation

**DOI:** 10.1101/2025.06.15.659604

**Authors:** Ruijiao Sun, Jason Cosens Walsman, Mark Wilber, Cheryl J. Briggs

## Abstract

Disease outcomes depend heavily on infection intensity which is often heterogeneous across and within host populations. Most individuals carry low pathogen loads and a few carry high loads, a pattern known as aggregation. While well-characterized in macroparasite systems, aggregation and infection intensity are rarely incorporated into microparasite models. This raises key questions: Do similar mechanisms underlie aggregation in macro- and microparasite systems? And how do aggregation and load-dependent effects shape outcomes such as host suppression and virulence-transmission trade-offs? To address these questions, we developed a novel continuous time microparasite model that allows the pathogen load distribution across hosts to evolve dynamically, shaped by within- and between-host processes. We applied this framework to the amphibian chytrid fungus system involving *Batrachochytrium dendrobatidis* (*Bd*), a fungal pathogen threatening amphibian populations worldwide. Our results show that load-dependent mortality reduces aggregation, while faster within-host replication increases it. Aggregation, in turn, weakens host suppression and flattens virulence-transmission trade-off, shifting peak transmission to higher replication rates. Overall, our continuous-time microparasite model provides new insights into how infection intensity and aggregation influence host-pathogen dynamics and offers a valuable framework for advancing theoretical and data-driven understanding of how within-host processes scale to population-level disease dynamics for microparasites.

## 1 Introduction

The outcome of infectious disease often depends on infection intensity of the host, as host survival (Wu, 2023), recovery (Wilber et al., 2016), and transmission potential (DiRenzo et al., 2014) are strongly influenced by the number of infectious agents carried by the host. Although the effects of infection intensity have primarily been studied in macroparasites both theoretically (Anderson & May, 1978, Fowler & Déirdre Hollingsworth, 2016, Grear & Hudson, 2011, Peacock et al., 2018) and empirically (Albery et al., 2021, Polak, 1996), infections caused by microparasites (viruses, bacteria, fungi, and protozoa) have mostly been treated as a binary variable, with individuals classified as either infected or uninfected (but see Walsman et al. (2024) and Wilber et al. (2021)). Macroparasite and microparasite models often represent opposite ends of the spectrum in terms of within-host processes. Macroparasite models typically assume that the parasites do not replicate on the hosts, whereas microparasite models frequently assume that within host replication is so rapid that following infection, the pathogen load immediately reaches a stable within-host carrying capacity. However, fluctuations in pathogen load over time can also be critical in microparasite systems for shaping host population dynamics and refining predictions of disease spread and severity.

Microparasite systems exhibit substantial variation in pathogen load within and among host populations, as well as under different environmental conditions, shaping host–pathogen dynamics. For instance, fatal H5N1 influenza cases correlate with elevated pharyngeal viral loads (de Jong et al., 2006). The bacterial pathogen *Mycoplasma gallisepticum* exhibits dynamic changes in load over the course of infection, influenced by the development of resistance and tolerance in host finch populations (Bonneaud et al., 2019). The fungal agent of White Nose Syndrome achieves very different pathogen loads in bats, across environmental conditions, within and between bat species (Langwig et al., 2015). Such findings underscore the need to integrate infection intensity into microparasite disease models, which exhibit within-host pathogen replication compared to macroparasites.

Following exposure to a pathogen, hosts may exhibit a continuum of outcomes, ranging from failed infections to carrying extremely high pathogen loads. Many hosts carry low pathogen loads, while few hosts carry high loads, a phenomenon known as “pathogen aggregation”. The degree of aggregation critically influences how parasites regulate host populations (Anderson, 1980), however, aggregation has been primarily considered for macroparasite, such as helminths and arthropods. Macroparasite aggregation typically follows a negative binomial distribution (Shaw & Dobson, 1995, Shaw et al., 1998), while microparasite infection loads quantified using molecular techniques is generally better represented by continuous distributions. Recent evidence suggests that microparasite show patterns of aggregation (Enriquez et al., 2022, Grogan et al., 2016, Schrock et al., 2025, Vale et al., 2013), indicating that aggregation levels may similarly regulate host-pathogen dynamics in microparasite diseases.

Many factors may drive pathogen aggregation. Studies of macroparasite systems reveal that host behavior (Johnson & Hoverman, 2014), variation in host susceptibility (Wilson et al., 2002), and pathogen virulence can shape aggregation patterns (Anderson & Gordon, 1982). Theoretical macroparasite models further suggest that reproduction directly on the host, a hallmark of microparasites, could amplify aggregation (Anderson & Gordon, 1982). Such within-host replication is rare in macroparasites compared to microparasite systems. Therefore, it is important to investigate whether the mechanisms maintaining aggregation and the implications of aggregation for host-parasite population dynamics are the same between macroparasites and microparasites.

Recent models have made progress in tracking microparasite pathogen load in populations using continuous distributions. Integral Projection Models (IPMs), which track heterogeneity in multiple continuously distributed variables within a population, have been adapted to monitor infection load in fungal pathogens (Walsman et al., 2024, Wilber et al., 2016, 2021). The reduced dimension IPMs, similar to early macroparsite models (Anderson & May, 1978, May & Anderson, 1978), allows for the distribution of pathogen loads to be represented by only a small number of moments which are statistical measures that describe key features of the distribution. Specifically, the first moment represents the mean, while the second moment captures the variance. Initial reduced dimension IPMs tracked only the mean of lognormally distributed load (Wilber et al., 2021), while a more recent work has extended these models to track both mean and variance (Walsman et al., 2024). While IPMs have provided valuable insight into load-dependent dynamics, they have not yet been applied to explore how changes in continuous pathogen load distributions influence host population dynamics which has important implications for community stability, conservation, and applied contexts such as pest control. A continuous-time modeling framework offers a promising next step, providing greater flexibility to incorporate additional mechanisms and to explore how dynamic pathogen distributions impact host–pathogen interactions.

Here, we develop ordinary differential equation (ODE) models to study the dynamics of pathogen load and aggregation patterns over continuous time. These ODE models are derived using the moment closure method, which reduces the dimensionality of a partial differential equation (PDE) model by approximating the full pathogen load distribution with a small number of moments. Specifically, moment closure allows us to track only the mean and variance of the pathogen load, assuming that the full distribution follows a lognormal shape. This approach is more computationally efficient and easier to implement than a full PDE model. We introduce two ODE models. Model 1 tracks only the first moment, dynamically modeling the mean pathogen load while assuming a fixed level of aggregation. This allows aggregation to be treated as a controllable parameter rather than an emergent property, enabling targeted exploration of its effects on disease outcomes. Model 2 tracks both the first and second moments of the PDE model, simultaneously capturing changes in the mean and variance of lognormally distributed pathogen loads.

We apply our model to a fungal pathogen disease system. Emerging fungal pathogens have posed significant threats to global biodiversity. Species such as *Batrachochytrium dendrobatidis, Ophidiomyces ophiodiicola*, and *Pseudogymnoascus destructans*, which infect amphibians, snakes, and bats, respectively, have led to population declines and extinctions in hundreds of wildlife species (Fisher & Garner, 2020, Hoyt et al., 2021, Lorch et al., 2016). Unlike most other microparasites characterized by short generation, high within-host replication rates, and lasting immunity, or macroparasites with longer generation and limited direct reproduction, fungal pathogens represent an intermediate case. We apply our models to a *Batrachochytrium dendrobatidis* (Bd), the fungus that causes chytridiomycosis and poses a global threat to amphibian populations (Fisher & Garner, 2020). Recent evidence suggest that *Bd* load distributions are consistently lognormal across diverse host species but with widely different levels of aggregation (Schrock et al., 2025). Using our models, we address three key questions: (Q1) What are the underlying mechanisms that are affecting pathogen aggregation levels? (Q2) How do pathogen load and aggregation affect host population dynamics? (Q3) What are the implications of pathogen load on the virulence-transmission trade-off? We used Model 2 with fluctuating variance to explore Q1, and Model 1 with dynamic mean pathogen load and fixed level of aggregation to explore Q2 and Q3.

## 2 Materials and Methods

### 2.1 A continuous-time model tracking pathogen load and load-dependent effects

#### 2.1.1 The full partial differential equation model

We propose a continuous-time disease model that captures the dynamics of continuous pathogen load distribution in infected hosts, incorporating within-host pathogen growth and load-dependent mortality. The model tracks three state variables: susceptible individuals, *S*(*t*); infected individuals with pathogen load *x, i*(*x, t*); and free-living environmental zoospores, *Z*(*t*) (Fig. 1). While the the model follows *Z*(*t*) on the natural scale, within-host pathogen load *x* is represented on the natural log scale, *x* ∈ (−∞, +∞), to account for the wide range of pathogen loads observed in empirical studies, which often span multiple orders of magnitude (Wu, 2023). The disease cycle (Fig. 1) begins when susceptible individuals encounter free-living zoospores in the environment, leading to infection with an initial pathogen load. Within-host pathogen growth drives changes in pathogen load, while host mortality rates depend on the pathogen load. Infected hosts shed zoospores into the environment, facilitating new infections. The full PDE model is presented below:

**Figure 1.**
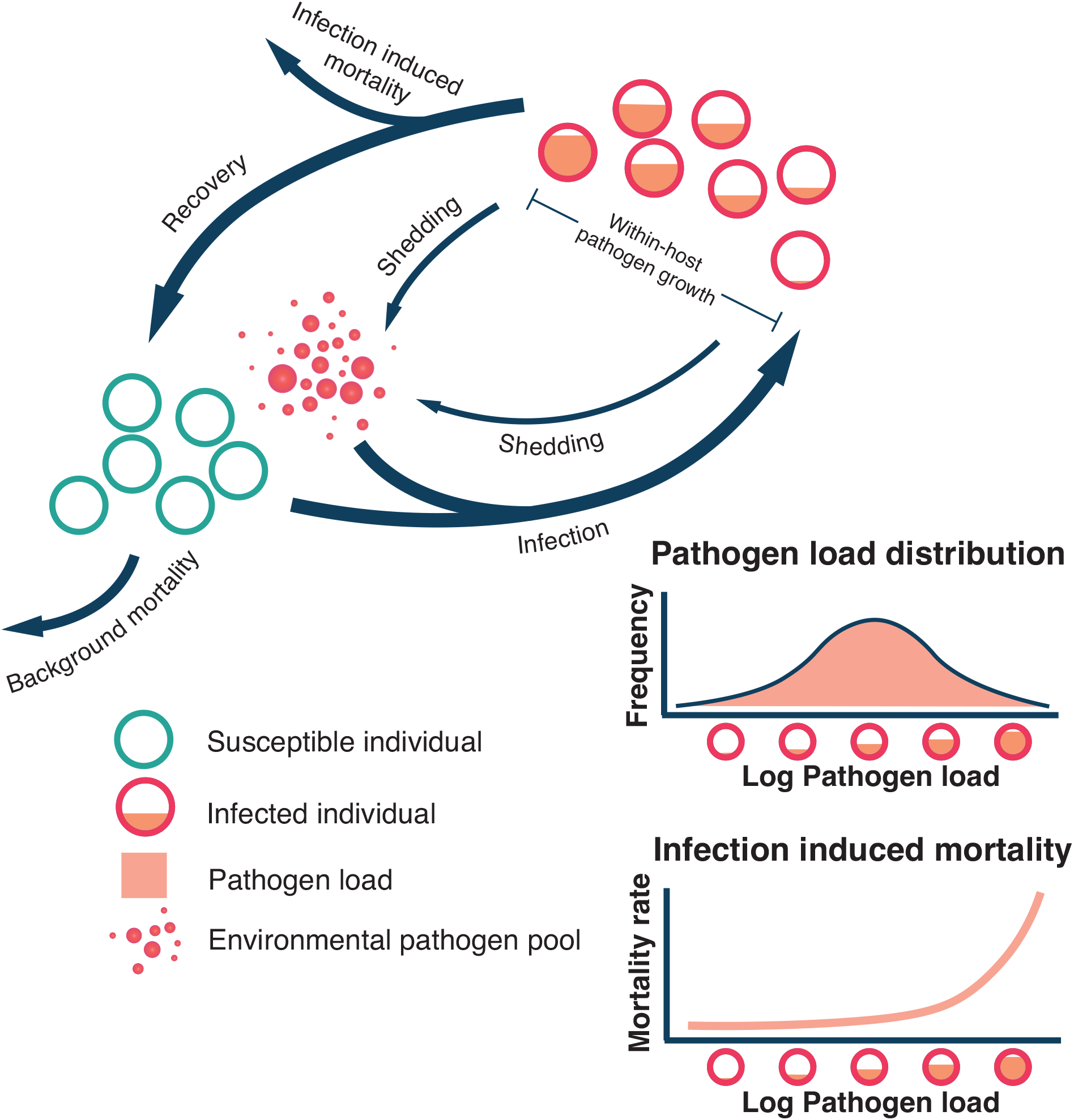
Schematic of the disease cycle. The infection begins with the encounter between pathogens and susceptible hosts, initiating within-host pathogen replication. Infected individuals may recover or die, and can continue shedding pathogens into the environment, leading to the infection of additional susceptible hosts. Mortality is load-dependent, with higher pathogen loads associated with an increased risk of death. Pathogen loads typically vary among infected hosts, often following a normal distribution on a log scale, with most individuals carrying low loads and a few carrying high loads.

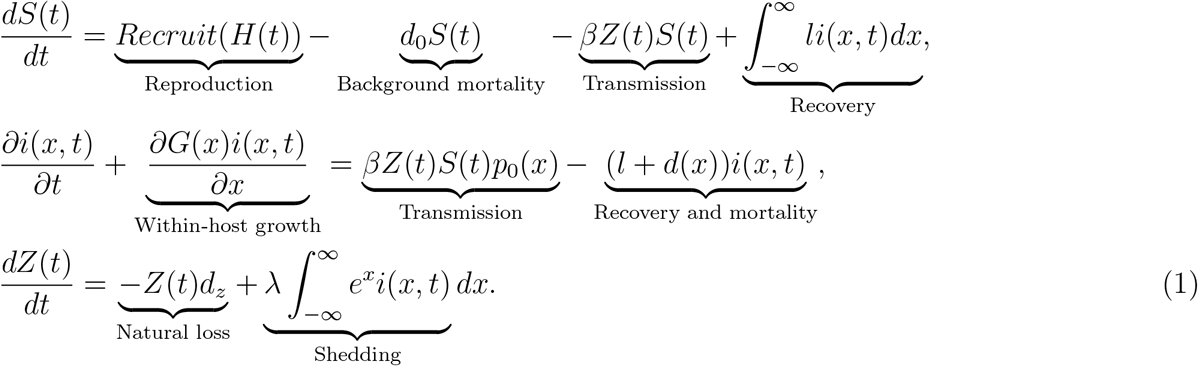

The dynamics of the susceptible population 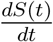 are influenced by recruitment from reproduction, recovery from infection, new infections, and background mortality. Recruitment is modeled as a density dependent function *Recruit*(*H*(*t*)) = *rH*(*t*)*e*^−*γH*(*t*)^, where 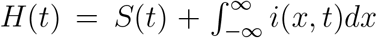 is the total host population, *r* is the maximum recruitment rate, *γ* is a parameter that determines the strength of the density dependence. The natural background instantaneous mortality rate is *d*_0_. *βZ*(*t*)*S*(*t*) is the new infection function and describes the rate at which susceptible host become infected, where *β* is the transmission coefficient. Recovery from infection is given by 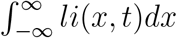, representing the total number of individuals recovering from infection across the pathogen load distribution. Here, we assume a constant recovery rate *l*, independent of pathogen load (load-dependent host recovery has been included in reduced-dimension IPMs (Walsman et al., 2024, Wilber et al., 2021).

We follow the Kermack and McKendrick epidemiology model in Kermack et al. (1997) to derive the changes in the infected population *i*(*x, t*) with respect to pathogen load *x* and time *t* by considering the conservation of individuals as they progress through the infection processes. The rate of change with respect to time *t* of infected individuals of load *x* would exactly balance the derivative with respect to load *x*, were it not for the force of recovery, mortality, and new infection. The pathogen-load of initial infection follows a normal distribution on a natural log scale with a probability density function of *p*_0_, with a mean of *µ*_0_ and standard deviation *σ*_0_.

Load-dependent mortality rate *d*(*x*) increases exponentially with the increase in pathogen load and is given by *d*(*x*) = *d*_0_ + *ab*^*x*^, where *d*_0_ is the baseline mortality rate, *a* (*a >* 0) and *b* (*b >* 1) determine how rapidly mortality increases with pathogen load, the severity of the pathogen’s influence on host mortality (Appendix Fig. S1). The rate of within-host pathogen growth is given by 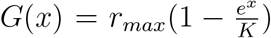 assuming logistic growth on the natural scale with a maximum growth rate of *r*_*max*_ and carrying capacity of *K*.

We assume that the pathogen shedding rate is proportional to the pathogen load with *λ* representing the shedding rate per unit pathogen load per unit time. The total amount of pathogen load in the population is measured by integrating over all infected individuals, where *e*^*x*^ converts log pathogen load to the natural scale. We also include the natural loss rate of environmental zoospores, denoted by *d*_*z*_.

We also explore the properties of the PDE model with diffusion term to simulate the behavior of host-parasite disease system with stochastic within-host pathogen growth as observed in many empirical study (Appendix Section S1.1).

#### 2.1.2 Model 1: First-order approximation with dynamic mean pathogen load and fixed aggregation by moment closure

Simplifying the PDE model to a set of ODE enhances both analytical tractability for gaining conceptual insights and computational efficiency. To achieve this, we use the moment closure method to develop a first-order approximation for the PDE model, which captures changes in the mean pathogen load while assuming a fixed variance in pathogen load. This approximation relies on the assumption that the pathogen load follows a lognormal distribution (Schrock et al., 2025). By manipulating the level of aggregation in the model, we can explicitly explore the effects of pathogen load aggregation on the outcomes of infectious disease dynamics (Q2 and Q3).

Here, rather than tracking the full distribution of log pathogen load *x* for infected population *i*(*x, t*), we track the first moment of *x*, i.e., the mean pathogen load. We track four key state variables: *S*(*t*), the population of susceptible individuals; *I*(*t*), the total infected population given by 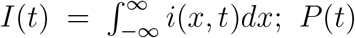, the total pathogen load in the infected population calculated as 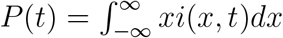 on a natural log scale; *Z*(*t*), the free-living zoospores in the environments on the natural scale. To approximate the pathogen load distribution, we use a probability density function for lognormally distributed pathogen load with a mean 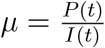 and a fixed standard deviation *σ*_*F*_. The final form of the first order approximation is:

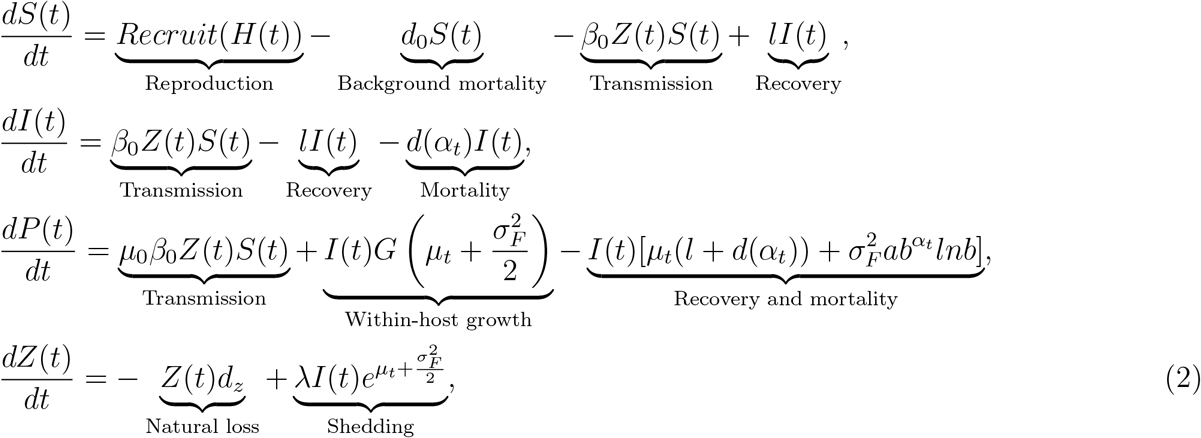

where 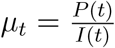 and 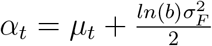. Detailed derivation is presented in Appendix Section

Macroparasite models frequently assume that the distribution of parasite across hosts follow a negative binomial distribution, in which the degree of aggregation is determined by the dispersion parameter *k*, with smaller *k* means higher level of aggregation. Aggregation occurs when the variance is greater than the mean. In our microparasite model, we assume that the distribution of the pathogen across hosts is best represented by a log-normal distribution, where aggregation is determined by the standard deviation *σ* of the distribution. A higher *σ* corresponds to a higher level of aggregation. In this Model 1 with fixed aggregation level, we change the fixed *σ*_*F*_ to explore different aggregation scenarios. When both the mean and aggregation level are allowed to change, as in the full PDE model (Section 2.1.1) and the second-order approximation (Section 2.1.3), we quantify aggregation using the coefficient of variation (*CV*), defined as 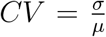, which represents the relative standard deviation of pathogen load. Higher *CV* values indicate greater aggregation.

#### 2.1.3 Model 2: Second-order approximation with dynamic mean pathogen load and changing aggregation by moment closure

To investigate the mechanisms contributing to pathogen aggregation (Q1), we also derive a second-order approximation of our PDE model, which tracks both the mean and variance of pathogen load under the assumption of a lognormal distribution. The second moment of pathogen load is defined as: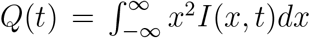. The variance of the lognormally distributed pathogen load, 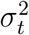, is given by: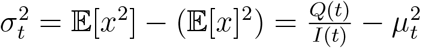. The final form of the second order approximation is:

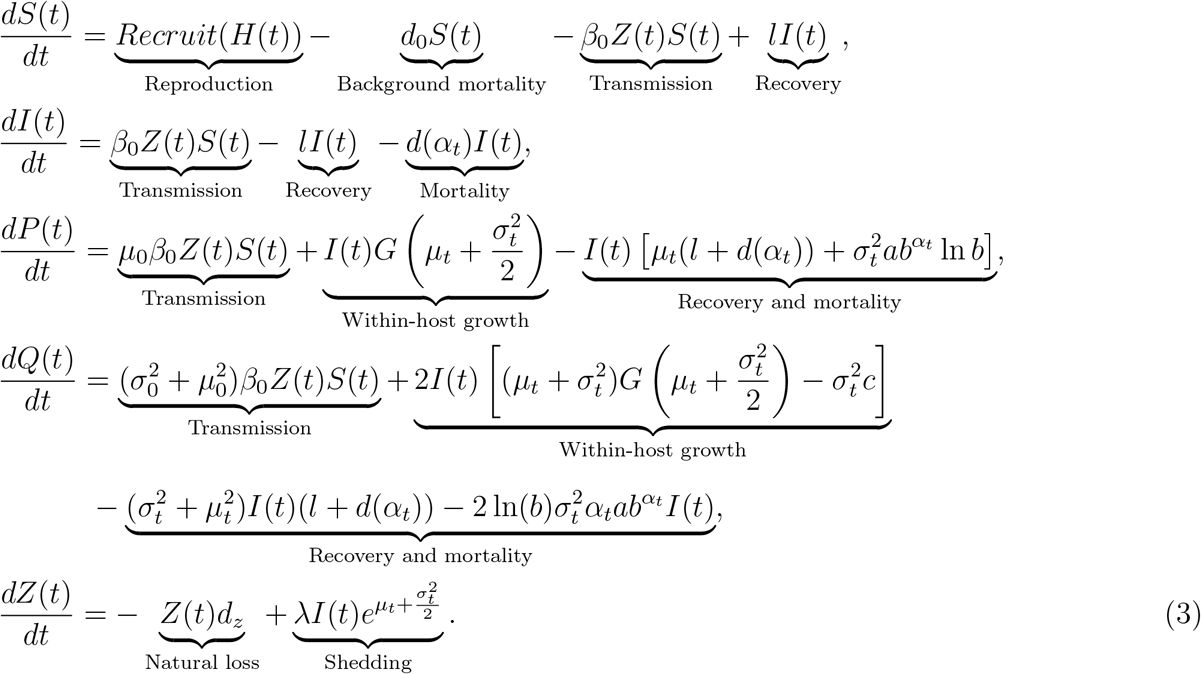

The mean of pathogen load is given by 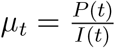 and the variance is 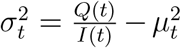. Detailed derivation is presented in Appendix Section S1.3.

#### 2.1.4 Connection to load-independent microparasite models

This framework remains consistent with classic load-independent microparasite models. Our model can be reduced to a load-independent structure when the within-host pathogen growth rate becomes infinitely fast (*r*_*max*_ → ∞) such that pathogen load reaches its carrying capacity almost instantaneously after infection. This eliminates within-host growth dynamics and variability, and the initial infection load approaches equilibrium load. As aggregation disappears (*σ* → 0), the pathogen load distribution becomes tightly concentrated around the equilibrium load, and all infected individuals carry approximately the same load *µ*. As a result, the mortality rate becomes independent of pathogen load, since all individuals are equally affected. Under these limiting conditions, pathogen load is effectively constant across hosts, and infection status alone determines transmission and mortality, consistent with the assumptions of standard load-independent microparasite models.

### 2.2 Application to the amphibian chytrid fungal disease system

To demonstrate the usefulness of our approach and explore pathogen-host dynamics in fungal diseases, we apply our model to the amphibian chytrid fungal disease system. *Batrachochytrium dendrobatidis* (*Bd*) has caused severe declines and extinctions in many amphibian species worldwide (Fisher & Garner, 2020, Luedtke et al., 2023, Scheele et al., 2019). These declines are strongly influenced by pathogen load and load-dependent mortality in infected frogs (Vredenburg et al., 2010). We parameterize our demographic parameters according to the California red-legged frog (*Rana draytonii*), which is threatened by *Bd*, faces additional challenges due to its endangered status (Russell et al., 2019). Because limited experimental data on *Bd* exist for this species, we draw disease parameters from the closely related mountain yellow-legged frog (*Rana mucosa*), which has similarly experienced declines driven by *Bd* and other stressors (Briggs et al., 2005, 2010). Detailed parameter values are provided in Appendix Table S1. Using these biologically informed parameters, we apply our modeling framework to test whether mechanisms driving aggregation in macroparasite systems also apply to microparasites, and to examine how aggregation and load-dependent effects shape host population dynamics and the virulence–transmission trade-off.

## 3. Results

### 3.1 Analytical insights

The equilibrium mean pathogen load, *µ*\*, and variance, (*σ*\*)^2^, of a lognormally distributed pathogen load are achieved at time *t* when:

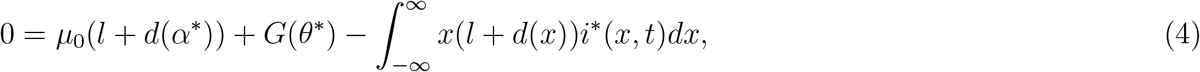

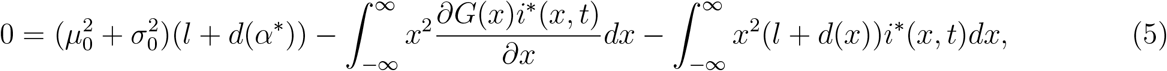

where 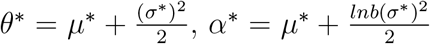, and *i*\*(*x, t*) represents the density function of the pathogen load distribution at equilibrium, which follows a normal distribution with mean *µ*\* and standard deviation *σ*\*. The equilibrium of the first moment (i.e., mean pathogen load, Eq. 4) and second moment of pathogen load (Eq. 5) reflects the balance between new infections and pathogen growth within the host. The equilibrium equation for mean pathogen load (Eq. 4) represents a weighted average of pathogen load contributions from within-host growth and initial infection, where the relative contributions depend on mortality (*d*(*x*)) and loss of infection (*l*). When loss of infection and mortality are high, the mean pathogen load approximates the initial infection load *µ*_0_. However, as loss of infection and mortality decrease, the equilibrium mean pathogen load approaches pathogen carrying capacity on the host. Similarly, the equilibrium of the second moment (Eq. 5) reflects a balance between variance contributions from initial infection and pathogen growth, offset by reductions due to recovery and mortality.

### 3.2 Load-dependent mortality and within-host pathogen growth affecting aggregation

We used Model 2, which incorporates both changing mean and variance in the lognormal distribution of pathogen load, to explore the factors influencing pathogen aggregation (Fig. 2, see results for full PDE model and Model 2 in Appendix Section S3). Our results show that both load-dependent mortality and within-host pathogen growth affect pathogen aggregation in the infected population (Fig. 2). The rate at which mortality increases with pathogen load varies depending on pathogen pathogenicity and host tolerance. Here, we interpret host tolerance as the degree to which mortality increases with rising pathogen load. Hosts with lower tolerance (higher *b* value) exhibit a steeper increase in mortality as pathogen load rises, whereas more tolerant hosts (lower *b* value) maintain lower mortality even at higher pathogen loads, resulting in a flatter load-dependent mortality curve (Appendix Fig. S1). As host tolerance decreases (i.e., the load-dependent mortality curve steepens), the coefficient of variation in pathogen load also decreases (Fig. 2a). This occurs because lower tolerance leads to fewer hosts with high pathogen loads, as these individuals experience high mortality(Fig. 2b). Consequently, the distribution shifts toward the initial pathogen load, with more individuals having similar, lower loads. In contrast, when host tolerance increases, hosts can survive with relatively higher pathogen loads, leading to a broader distribution with heavier tails(Fig. 2b).

**Figure 2.**
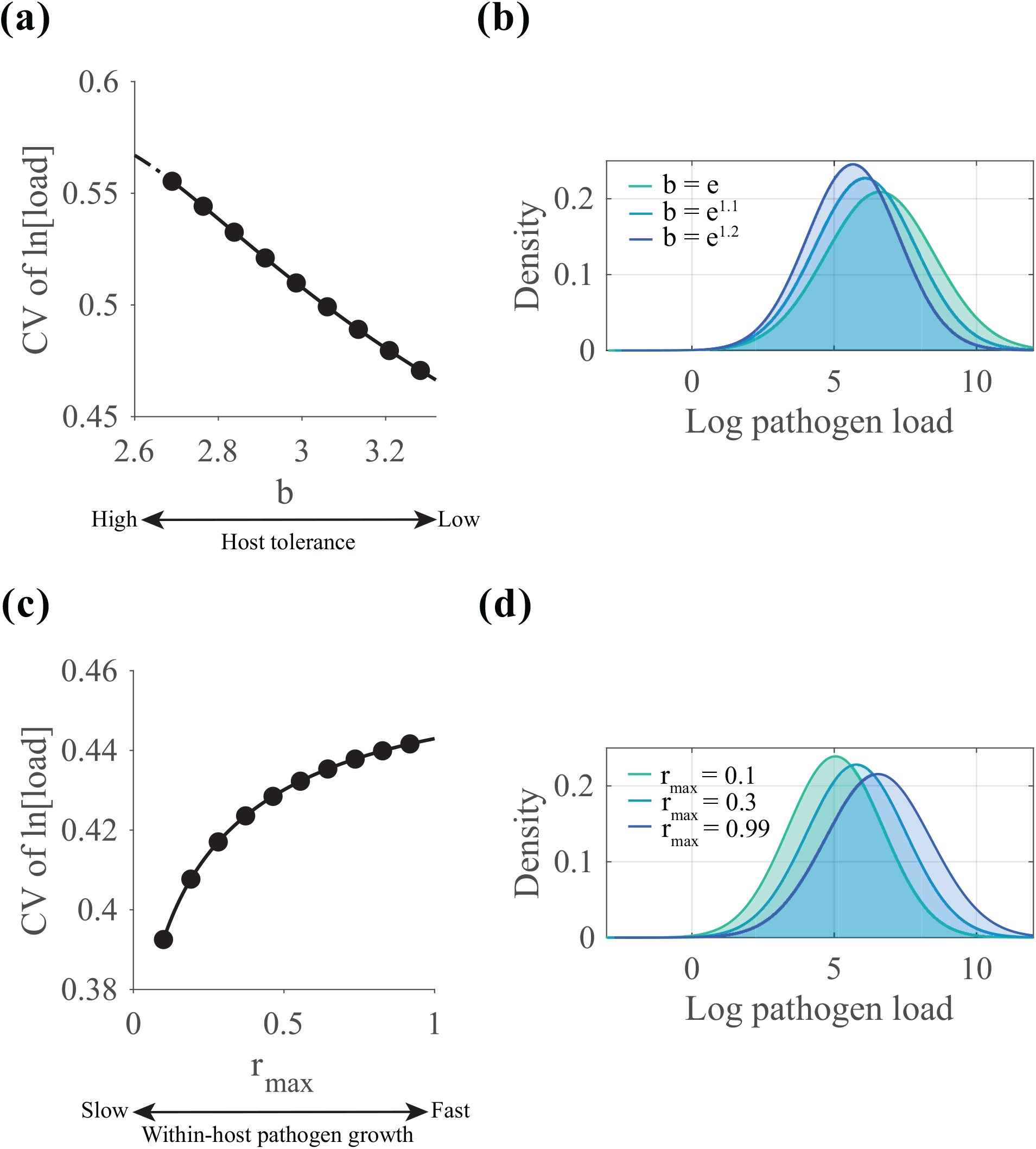
Infection processes influencing pathogen aggregation. (a) the steepness of load-dependent mortality (reflecting host tolerance) affects the aggregation of pathogen load, measured by the coefficient of variation [CV]; (b) pathogen load distributions under varying levels of load-dependent mortality; (c) the rate of within-host pathogen growth influences aggregation; (d) pathogen load distributions under different within-host growth rates.

Within-host pathogen growth also influences aggregation, with faster growth increasing the coefficient of variation (i.e., increasing aggregation) (Fig. 2c). This occurs because rapid within-host pathogen replication allows hosts to reach higher pathogen loads in a shorter time without dying. In contrast, when pathogen replication is slower, hosts are less likely to reach high loads, resulting in a lower mean pathogen load across the infected population and a more uniform distribution and lower levels of aggregation of pathogen loads (Fig. 2d).

### 3.3 Host population dynamics

We employ Model 1 with a changing mean and fixed variance 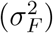 in the lognormal distribution of pathogen load, to investigate how pathogen aggregation, load-dependent mortality, and within-host replication affect host suppression (Fig. 3, see Appendix Section S3 for analysis using the full PDE model and Model 2). We show that as pathogen load aggregation increases (i.e., variance increases), the pathogen’s ability to suppress the host population declines (Fig. 2a) because heavily infected individuals are removed from the population before contributing significantly to onward transmission. Correspondingly, the basic reproductive number (*R*_0_) decreases with increased pathogen aggregation (Fig. 3c).

**Figure 3.**
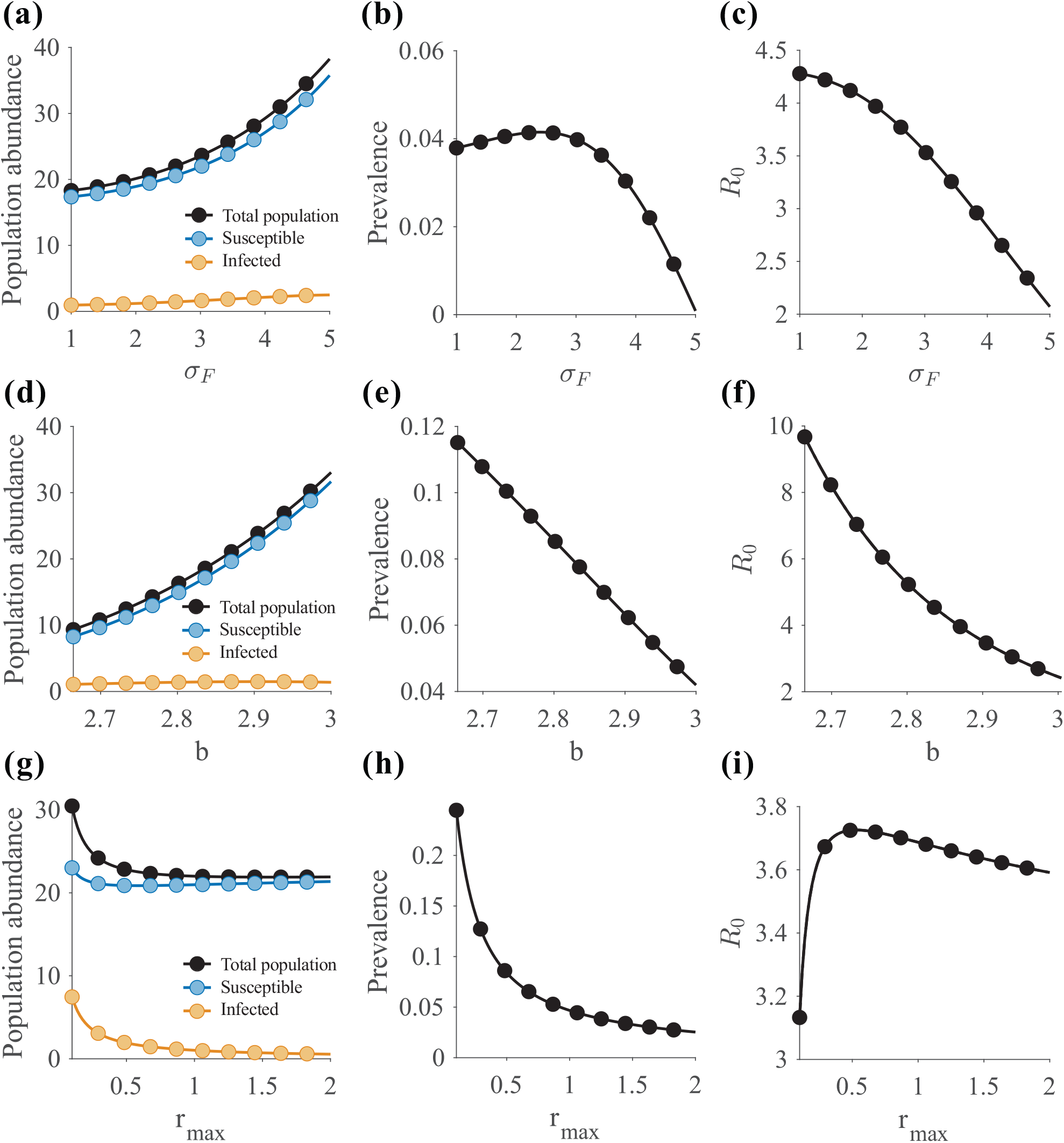
Effects of infection processes on host population dynamics and disease transmission potential (*R*_0_). Panels show the effects of (a–c) pathogen aggregation, (d–f) load-dependent mortality, and (g–i) within-host pathogen growth on host population suppression, disease prevalence, and the basic reproduction number (*R*_0_).

An interesting results of our model is the breakdown of the relationship between *R*_0_ and prevalence when infection intensity and aggregation are incorporated. In traditional Susceptible-Infected-Recovered (SIR) load-independent model, equilibrium prevalence and *R*_0_ are correlated (Diekmann et al., 1990). In our model, with increasing pathogen aggregation (higher *σ*_*F*_), prevalence initially increases even as mean pathogen load declines due to the selective removal of heavily infected hosts via load-dependent mortality (Fig. 3b and Appendix Fig. S5). This occurs because the transmission driven by shedding increases initially, dominated by the variance component of the load distribution. As a result, the infected population carries lower average pathogen loads, yet contributes to a larger environmental zoospore pool, leading to a rise in prevalence. Eventually, at high level of aggregation, this trend reverses as elevated mortality severely reduces the infected host pool, causing prevalence to decline. Therefore, even as *R*_0_ decreases, prevalence can initially rise due to increased shedding, then decline as mortality becomes the dominant force. When load-dependent mortality steepens, this initial increase in prevalence disappears because the effects of mortality dominate as aggregation increases (Appendix Fig. S6).

Furthermore, reduced host tolerance (higher *b* value and a steeper load-dependent mortality curve), leads to reduced host suppression effects (Fig. 3d), accompanied by decreased prevalence (Fig. 3e) and *R*_0_ (Fig. 3f) due to increased mortality among infected individuals. Faster pathogen replication rates enhance host suppression effects (Fig. 3g). Prevalence decreases with increasing within-host pathogen growth (Fig. 3h), likely because individuals rapidly reach higher pathogen loads, leading to higher mortality rates of infected individuals. *R*_0_ exhibits a virulence-transmission trade-off, maximized at an intermediate within-host pathogen growth rate (Fig. 3i).

### 3.4 Virulence transmission trade-offs

We also used Model 1 to explore virulence transmission trade-offs (Fig. 4, see results for full PDE model and Model 2 in Appendix Section S3). The virulence-transmission trade-off occurs only when the rate at which mortality increases with pathogen load exceeds the within-host pathogen growth. Otherwise, the effects of pathogen replication remains dominant, leading to consistently higher transmission with increasing replication rates, and no virulence-transmission trade-off occurs. Since our model assumes logistic within-host pathogen growth, we explore virulence-transmission trade-offs for *b* values greater than *e*.

**Figure 4.**
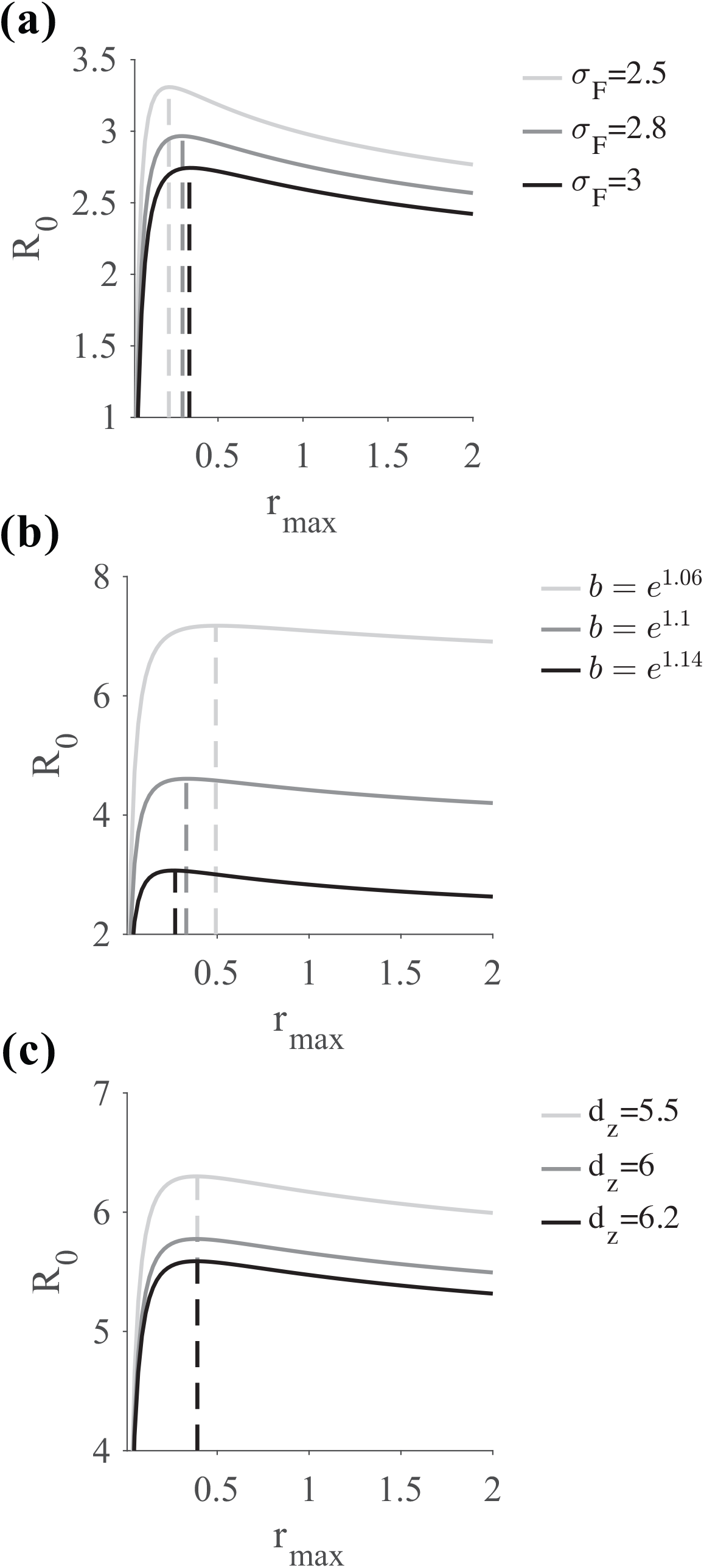
Factors influencing virulence–transmission trade-offs. Pathogen aggregation (a), load-dependent mortality (b), and the loss rate of environmental zoospores (c) shape the relationship between virulence and transmission. The dashed line indicates the level of within-host pathogen growth that maximizes the basic reproduction number (*R*_0_).

We find that increased pathogen aggregation dampens the virulence-transmission trade-off curve and reduces the maximum *R*_0_ (Fig. 4a), as mean pathogen load decreases with higher aggregation (Appendix Fig. S5). *R*_0_ is maximized at a higher within-host pathogen growth rate under greater aggregation because because replication does not uniformly raise individual pathogen loads, and the associated mortality costs are delayed. When host tolerance decreases (i.e., higher *b* values and a steeper load-dependent mortality curve), the maximum *R*_0_ also decreases, but the trade-off curve becomes more pronounced (Fig. 4b). *R*_0_ is then maximized at a lower within-host pathogen growth rate, as a lower growth rate is needed to offset the increased mortality of infected individuals. We also examined the effects of environmental zoospore loss rates (*d*_*z*_) on virulence-transmission trade-offs (Fig. 4c). Higher environmental loss rates reduce the maximum *R*_0_. Our model assumes that infection occurs exclusively from the environmental zoospore pool, with no direct transmission from infected individuals, so changes in environmental persistence do not impact the evolution of virulence. Consequently, the equilibrium mean pathogen load remains independent of the environmental loss rate, and *R*_0_ is always maximized at the same pathogen growth rate.

## 4 Discussion

Infection intensity is a key source of individual infection heterogeneity, but is rarely considered in microparasite models. Recent evidence suggests that aggregation levels vary across populations (Schrock et al., 2025), and both within-host and between-host process may contribute to different aggregation levels. Progress has been made in addressing theoretical gaps using IPM models, particularly in understanding pathogen invasion and virulence evolution (Walsman et al., 2024, Wilber et al., 2021). To further investigate the factors driving aggregation and host suppression, we develop a continuous-time model that is both analytically tractable and easy to implement. Our model accounts for: (1) Pathogen load-dependent mortality rates across a range of function behaviors, capturing different levels of host tolerance or virulence; (2) Logistic within-host pathogen growth; (3) A continuous pathogen load distribution, approximated using a lognormal distribution that is both mathematically valid and ecologically meaningful. Using the amphibian-chytrid fungal disease system as a case study, our results align closely with findings from macroparasite models. This highlights the importance of incorporating pathogen load and infection intensity heterogeneity into microparasite models to accurately infer host-pathogen dynamics.

Different levels of pathogen load aggregation have been documented across many microparasite systems, both between populations of different host species and within the same host population under varying environmental conditions (Enriquez et al., 2022, Schrock et al., 2025, Vale et al., 2013). Consistent with theoretical predictions from macroparasite models (Anderson & Gordon, 1982), our microparasite model predicts that stronger load-dependent mortality reduces pathogen aggregation, while faster within-host pathogen replication increases aggregation. Empirical evidence from microparasite systems supports these predictions. In the *Daphnia magna* and the bacterial parasite *Pasteuria ramosa* systems, improved environmental conditions, which increase host tolerance and reduce pathogen virulence (Vale et al., 2011), are associated with higher levels of parasite load aggregation (Vale et al., 2013). These findings reveal how host-pathogen interactions shape aggregation patterns, and how environmental factors influencing these processes can also play a key role in determining aggregation levels. Thus, aggregation patterns can offer insights into underlying epidemiological processes driving changes in transmission potential and virulence evolution.

Our findings align with theoretical predictions from macroparasite models (Anderson, 1980), emphasizing the key roles of pathogen aggregation, load-dependent mortality, and withinhost pathogen growth in shaping host suppression effects in microparasite diseases (Fig. 3). These dynamics may help explain patterns of host–pathogen coexistence in endemic systems and are supported by empirical observations. For instance, in a recovered amphibian community affected by *Bd*, Fleay’s barred frog (*Mixophyes fleayi*) exhibits rare instances of high pathogen loads and lower prevalence compared to sympatric species, underscoring the role of aggregation in modulating endemic disease patterns (Hollanders et al., 2023). Variation in tolerance and resistance also explain divergent population trajectories (Fig. 2). For example, boreal toads (*Anaxyrus boreas boreas*) have suffered severe declines in Colorado due to *Bd*, whereas populations in Wyoming appear less impacted, likely due to higher tolerance and slower within-host Bd growth (Hardy et al., 2025). The Wyoming population also exhibits higher prevalence level (Muths et al., 2011), which aligns with our model predictions. Given the strong interactions among load-dependent mortality, within-host pathogen growth, and aggregation as suggested in our results, integrating these factors into microparasite models is essential for predicting population-level outcomes and understanding why some host populations persist despite chronic infections.

Empirical evidence from macroparasite systems generally shows that higher levels of aggregation are associated with lower prevalence (Shaw & Dobson, 1995, Shaw et al., 1998). When we assume a load-dependent mortality function that increases slightly faster than exponentially with log pathogen load in our microparasite model, we find that prevalence initially rises at low aggregation levels but subsequently declines as aggregation continues to increase, driven by the non-linearity of load-dependent mortality on the log scale (Fig. 3b). When load-dependent mortality is even steeper, prevalence decreases monotonically with increasing aggregation (Appendix Fig. S6). Thus, realistic load-dependent mortality relationships may be considerably steeper than linear on the natural scale and may represent a widespread feature of host–parasite systems, as supported by empirical studies (Gupta & Vale, 2017, Louie et al., 2016, Wu, 2023).

Strong pathogen aggregation dampens the virulence-transmission trade-off, consistent with the findings in Wilber et al. (2021). Although virulence–transmission trade-offs have been widely predicted in theoretical studies, empirical support has been limited (Acevedo et al., 2019, Ebert & Bull, 2003). This is potentially because changes in aggregation influence the emergence of these trade-offs, while most previous studies have assumed homogeneous pathogen loads (Alizon & van Baalen, 2005, Day, 2002). We show that as the pathogen aggregation increases, *R*_0_ is maximized at higher within-host pathogen growth rates, suggesting that aggregation may select for increased pathogen virulence. Previous research has primarily focused on static heterogeneity in host populations, with mixed predictions on its impact on virulence evolution (Ganusov et al., 2002, Osnas & Dobson, 2012). For example, random heterogeneity in demographic parameters has been shown to increase virulence with rising heterogeneity (Ganusov et al., 2002), while heterogeneity involving trade-offs in parasite fitness predicts reduced virulence (Ebert & Mangin, 1997, Regoes et al., 2000). However, dynamic heterogeneity, such as variability in pathogen load resulting from factors not fixed for a specific host has received less attention. Our results demonstrate that dynamic heterogeneity can dampen the strength of virulence transmission trade-offs while also altering the overall value of *R*_0_. Given that variability in infection intensity is nearly universal across infections, incorporating this dynamic heterogeneity is crucial for accurately prediction for the evolution of virulence.

Our results also show that virulence-transmission trade-offs depend on mortality accelerating faster with load than transmission does, aligning with previous theoretical predictions assuming a homogeneous pathogen load (Alizon & van Baalen, 2005, Day, 2002). Since we assume transmission is proportional to host pathogen load on the natural scale, the trade-off is dampened when mortality increases less steeply with pathogen load (i.e., when *b* is larger but close to *e*) even when load-dependent mortality is present. Unlike the results of the IPM model in Wilber et al. (2021), where steeper declines in survival with increasing load lead to higher maximum *R*_0_, our model predicts lower maximum *R*_0_ under steeper mortality functions. This difference arises because Wilber et al. (2021) alters the shape of the survival curve directly in a way that also increases survival for individuals with lower pathogen loads when the survival curve is steeper. Our findings emphasize that *R*_0_ is highly sensitive to the functional form of load-dependent mortality, highlighting the importance of using biologically realistic functions. In our model, we assume transmission increases proportionally with pathogen load on the natural scale. However, in real systems, factors such as host behavior and physiology can modify this relationship (VanderWaal & Ezenwa, 2016), potentially affecting the occurrence of virulence-transmission trade-offs. Despite strong theoretical support for the virulence-transmission trade-off, empirical evidence for this relationship remains limited (Froissart et al., 2010). Therefore, measuring the steepness of load-dependent mortality curves and how transmission potential changes with pathogen load could help identify host-parasite systems where virulence-transmission trade-offs are more likely to occur.

Our results reveal a complex interplay between load-dependent mortality and within-host replication in shaping population outcomes and disease prevalence. Most previous models have treated microparasites and macroparasites as distinct, emphasizing within-host multiplication for microparasites and external reproduction for macroparasites (Anderson, 1979). This has often led to pathogen load being tracked only in macroparasite models, while microparasite models only track infection status. However, this separation may oversimplify the continuum of parasite life-history strategies observed in nature. Microparasite replication rates may vary widely across genotype of strains (Little et al., 2008), host populations (Hardy et al., 2025), and environmental conditions (Vale et al., 2011). Moreover, some macroparasites also exhibit direct replication on hosts (Stepek et al., 2006). Thus, there is a clear need for integrative models that bridge the microparasite–macroparasite continuum by incorporating pathogen aggregation, load-dependent effects, and within-host dynamics to improve our understanding of host–parasite interactions and their demographic consequences for host species.

Our continuous-time microparasite model offers a computationally efficient, empirically relevant, and analytically tractable framework for advancing our understanding of pathogen load dynamics. By incorporating load-dependent effects and tracking changes in the distribution of pathogen loads among infected individuals, our results demonstrate that both pathogen load distributions critically influence host suppression, disease prevalence, and invasion potential in microparasite systems. These findings highlight the importance of integrating load-dependent processes and infection intensity heterogeneity into both theoretical and empirical studies of microparasite. More broadly, our model provides a foundation for datadriven inference using population-level observations and offers a flexible tool for exploring host–parasite dynamics across a wide range of ecological contexts.

## 5 Acknowledgement

This work was supported by the University of California Climate Action 2023 Seed Award R02CP7252 to R.S. and C.B.; Hatch Project 7001607 to M.W.; and National Science Foundation grant DBI-2120084 to J.C.W., M.W., and C.B.

## 6 Author contribution

R.S.: conceptualization, methodology, investigation, interpretation, writing original draft;

J.W.: methodology, interpretation, writing and editing;

M.W.: methodology, interpretation, funding acquisition, writing and editing;

C.B.: conceptualization, methodology, interpretation, funding acquisition, project administration, writing and editing.

All authors provided final approval for publication, and accepted responsibility for the work conducted.

## Notes

### Competing Interest Statement

The authors have declared no competing interest.

