## Appendix S1 for "A continuous-time microparasite model incorporating infection intensity and parasite aggregation"

### Section S1 Model derivation and approximation using moment closure

#### Section S1.1 Partial differential equation (PDE) model

The full partial differential equation (PDE) model tracks three state variables $S(t)$ - susceptible individuals, $i(x,t)$ - infected individuals with pathogen load $x$, and $Z(t)$ - environmental zoospores. The initial distribution of pathogen load among newly infected hosts follows a probability density function $f_0(x)$. The full PDE model with deterministic within-host pathogen growth is presented below:

$$\frac{dS(t)}{dt} = \underbrace{Recruit(H(t))}_{\text{Reproduction}} - \underbrace{d_0 S(t)}_{\text{Background mortality}} - \underbrace{\beta Z(t)S(t)}_{\text{Transmission}} + \underbrace{\int_{-\infty}^{\infty} li(x,t)dx}_{\text{Recovery}},$$

$$\frac{\partial i(x,t)}{\partial t} + \underbrace{\frac{\partial G(x)i(x,t)}{\partial x}}_{\text{Pathogen growth}} = \underbrace{\beta Z(t)S(t)f_0(x)}_{\text{Transmission}} - \underbrace{(l+d(x))i(x,t)}_{\text{Recovery and mortality}},$$

$$\frac{dZ(t)}{dt} = \underbrace{-Z(t)d_z}_{\text{Natural loss}} + \underbrace{\lambda\int_{-\infty}^{\infty} e^x i(x,t)\,dx}_{\text{Shedding}}.$$

Many pathogens exhibit stochastic growth patterns. To capture this, we also derived a full PDE model incorporating stochastic within-host pathogen growth, represented by a diffusion term $D = \frac{\sigma_G^2}{2}$, where $\sigma_G$ denotes the standard deviation of the within-host pathogen growth rate. The dynamics of infected individuals are then described by:

$$\frac{\partial i(x,t)}{\partial t} + \underbrace{\frac{\partial G(x)i(x,t)}{\partial x}}_{\text{Pathogen growth}} = \underbrace{\beta Z(t)S(t)f_0(x)}_{\text{Transmission}} - \underbrace{(l+d(x))i(x,t)}_{\text{Recovery and mortality}} + \underbrace{D\frac{\partial^2 i(x,t)}{\partial x^2}}_{\text{Diffusion}}. \tag{S1}$$

The individual pathogen load on log scale ($x$) can change due to within-host pathogen growth, and the growth rate is given by $\frac{dx}{dt} = G(x)$. We model within-host pathogen growth following logistic growth on a natural scale, and the growth of pathogen load on natural scale $(X)$ is given by $\frac{dX}{dt} = r_{max}(1 - \frac{X}{K})X$. Then, the growth function $G(x)$ on log scale is derived as follows:

$$G(x) = \frac{dx}{dt} = \frac{d(lnX)}{dt} = \frac{d(lnX)}{dX}\frac{dX}{dt} = \frac{1}{X}\frac{dX}{dt} = r_{max}(1 - \frac{X}{K}) = r_{max}(1 - \frac{e^x}{K}).$$

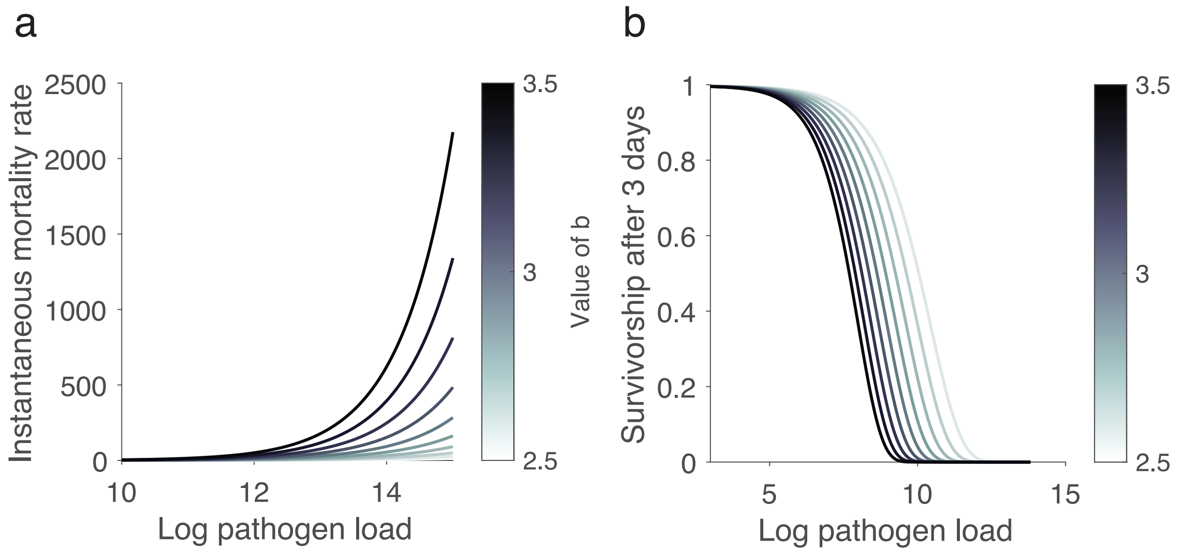


Figure S1: Load-dependent mortality as a function of pathogen load, showing variation in the severity of pathogen impact on the host. Panels show the effects of different values of $b$: (a) instantaneous mortality rate, and (b) three-day survival probability.

Load-dependent mortality rate $d(x)$ increases exponentially as the increase of pathogen load and is given by $d(x) = d_0 + ab^x$, where $d_0$ is the baseline mortality rate, $a$ $(a > 0)$ and $b$ $(b > 1)$ determine how rapidly mortality increases with pathogen load, reflecting the severity of the pathogen's influence on host mortality (Fig. S1a). As pathogen load increases, the three-day survival probability declines sharply, with steeper declines observed for larger values of $b$ (Fig. S1b).

#### Section S1.2 1st order approximation

We develop a first-order approximation of the deterministic PDE model using the moment closure method, assuming that pathogen load follows a log-normal distribution with probability density function $f(x; \mu_t, \sigma_F)$. This resulting ordinary differential equation (ODE) model tracks the changes in mean pathogen load $(\mu)$, while assuming a fixed standard deviation $(\sigma_F)$. We also assume the initial infection follows a log-normal distribution with a probability density function $f_0(x; \mu_0, \sigma_0)$, characterized by a mean of $\mu_0$ and a standard deviation of $\sigma_0$. We tracked four state variable $S(t)$ - population of susceptible individuals, and $I(t) = \int_{-\infty}^{\infty} i(x,t)dx$ - total population of infected individuals, $P(t)$ - total pathogen load of the infected population, $Z(t)$ - free-living zoospores in the environments. And the mean pathogen load is given by $\mu = \frac{P(t)}{I(t)}$. The detailed derivations for the first-order approximation are presented in the following sessions.

##### Section S1.2.1 Model derivations

The the dynamics of susceptible individuals is then given by

$$\frac{dS(t)}{dt} = -d_0 S(t) - \beta Z(t)S(t) + lI(t) + R(N(t)).$$

For the infected population, the rate of change for the infected hosts with respect to time $t$ and pathogen load $x$ is given by

$$\frac{\partial i(x,t)}{\partial t} + \frac{\partial G(x)i(x,t)}{\partial x} = \underbrace{\beta Z(t)S(t)f_0(x;\mu_0,\sigma_0)}_{\text{Transmission}} - \underbrace{(l+d(x))i(x,t)}_{\text{Recovery and mortality}}.$$

Rearranging terms, we obtain

$$\frac{\partial i(x,t)}{\partial t} = \beta_0 Z(t)S(t)f_0(x;\mu_0,\sigma_0) - \frac{\partial G(x)i(x,t)}{\partial x} - (l+d(x))i(x,t).$$

We can then derive the rate of change in the total number of infected individuals $I(t)$ as follows:

$$\begin{aligned}\frac{dI(t)}{dt} &= \frac{d}{dt}(\int_{-\infty}^{\infty} i(x,t)dx)\\ &= \int_{-\infty}^{\infty} \frac{\partial i(x,t)}{\partial t}dx\\ &= \int_{-\infty}^{\infty} \beta_0 Z(t)S(t)f_0(x;\mu_0,\sigma_0)dx - \int_{-\infty}^{\infty}\frac{\partial G(x)i(x,t)}{\partial x}dx - \int_{-\infty}^{\infty}(l+d(x))i(x,t)dx.\end{aligned}$$

Since log pathogen load of infected population follows a normal distribution, $i(x,t)$ decays faster than an exponential function as pathogen load $x$ approaches $\pm\infty$. Therefore, we have

$$\int_{-\infty}^{\infty}\frac{\partial G(x)i(x,t)}{\partial x}dx = G(x)i(x,t)|_{-\infty}^{\infty} = \lim_{x\to\infty} G(x)i(x,t) - \lim_{x\to-\infty} G(x)i(x,t) = 0.$$

The total mortality of infected individuals is given by:

$$\begin{aligned}\int_{-\infty}^{\infty} d(x)i(x,t)dx &= I(t)\int_{-\infty}^{\infty} d(x)f(x;\mu_t,\sigma_F)dx\\ &= d_0 I(t) + I(t)\int_{-\infty}^{\infty}\frac{1}{\sigma_F\sqrt{2\pi}}e^{\frac{-(x-\mu_t)^2}{2\sigma_F^2}}ab^x dx\\ &= d_0 I(t) + I(t)ae^{(\mu_t+\frac{\sigma_F^2 lnb}{2})lnb}\\ &= d_0 I(t) + I(t)ab^{\alpha_t}\\ &= d(\alpha_t)I(t),\end{aligned}$$

where $\alpha_t = \mu_t + \frac{\sigma_F^2 lnb}{2} = \frac{P(t)}{I(t)} + \frac{\sigma_F^2 lnb}{2}$ and $f(x;\mu_t,\sigma_F)$ denotes the normal probability density function of log pathogen load $x$ with mean $\mu_t$ and standard deviation $\sigma_F$. Then, the final form for the dynamics of infected individuals is given by

$$\frac{dI(t)}{dt} = \beta_0 Z(t)S(t) - lI(t) - d(\alpha_t)I(t).$$

The changes in the total pathogen load of infected individuals $P(t) = \int_{-\infty}^{\infty} xi(x,t)dx$ include (1) pathogen loss due to load-dependent mortality, (2) host recovery, (3) within-host pathogen growth, and (4) the contribution of newly infected individuals with pathogen load $x$ with probability $p_0(x)$ from initial infection. The dynamics of $P(t)$ is given by

$$\begin{aligned}\frac{dP(t)}{dt} &= \frac{d}{dt}\int_{-\infty}^{\infty} xi(x,t)dx = \int_{-\infty}^{\infty} x\frac{\partial i(x,t)}{\partial t}dx \\ &= \underbrace{\int_{-\infty}^{\infty} x\beta_0 Z(t)S(t)f_0(x;\mu_0,\sigma_0)dx}_{\text{Transmission}} - \underbrace{\int_{-\infty}^{\infty} x\frac{\partial G(x)i(x,t)}{\partial x}dx}_{\text{Within-host pathogen growth}} - \underbrace{\int_{-\infty}^{\infty} x(l+d(x))i(x,t)dx}_{\text{Recovery and mortality}}.\end{aligned}$$

The pathogen loss through load-dependent mortality is given by:

$$\begin{aligned}\int_{-\infty}^{\infty} xd(x)i(x,t)dx &= I(t)\int_{-\infty}^{\infty} xd(x)f(x;\mu_t,\sigma_F)dx \\ &= I(t)\int_{-\infty}^{\infty} x(d_0+ab^x)f(x;\mu_t,\sigma_F)dx \\ &= I(t)\int_{-\infty}^{\infty} d_0 xf(x;\mu_t,\sigma_F)dx + I(t)\int_{-\infty}^{\infty} xab^x f(x;\mu_t,\sigma_F)dx \\ &= \mu_t d_0 I(t) + aI(t)\int_{-\infty}^{\infty} xb^x f(x;\mu_t,\sigma_F)dx.\end{aligned}$$

We use the moment-generating function to solve $\int_{-\infty}^{\infty} xb^x f(x;\mu_t,\sigma_F)dx$. We write the integral as $\int_{-\infty}^{\infty} xe^{xlnb} f(x;\mu_t,\sigma_F)dx$, which is the expectation of variable $xe^{xlnb}$ with a normal distribution, i.e., $\mathbb{E}(xe^{xlnb})$.
Based on the moment-generating function of normal distribution evaluated at $lnb$, we have

$$\mathbb{E}(e^{xlnb}) = e^{\mu lnb+\frac{\sigma_F^2(lnb)^2}{2}}$$

To derive $\mathbb{E}(xe^{xlnb})$, we took the derivative of $\mathbb{E}(e^{xlnb})$. Let $lnb = m$, we have $\mathbb{E}(e^{xlnb}) = \mathbb{E}(e^{mx})$, so that $\mathbb{E}(xe^{mx}) = \frac{d\mathbb{E}(e^{mx})}{dm}$. Then, we have

$$\mathbb{E}(xe^{xlnb}) = \mathbb{E}(xe^{mx}) = \frac{d\mathbb{E}(e^{mx})}{dm} = \frac{de^{\mu_t m+\frac{\sigma_F^2(m)^2}{2}}}{dm} = (\mu+\sigma_F^2 m)e^{\mu_t m+\frac{\sigma_F^2(m)^2}{2}}.$$

Substituting $m = lnb$ back, we obtain

$$\int_{-\infty}^{\infty} xe^{xlnb}i(x,t)dx = \mathbb{E}(xe^{xlnb}) = (\mu_t+\sigma_F^2 lnb)e^{\mu_t lnb+\frac{\sigma_F^2(lnb)^2}{2}}.$$

Combining both components of the integral, the total loss of pathogen load due to mortality is given by:

$$\begin{aligned}\int_{-\infty}^{\infty} xd(x)i(x,t)dx &= \mu_t d_0 I(t) + aI(t)(\mu_t+\sigma_F^2 lnb)e^{\mu_t lnb+\frac{\sigma_F^2(lnb)^2}{2}} \\ &= I(t)[\mu_t d(\alpha_t) + \sigma_F^2 lnbab^{\alpha_t}].\end{aligned}$$

The pathogen load loss due to the recovery of infected individuals is $\int_{-\infty}^{\infty} xli(x,t)dx = l\mu_t I(t)$.

For the pathogen load change from within-host pathogen growth, we use integration by parts:

$$\int_{-\infty}^{\infty} x\frac{\partial G(x)i(x,t)}{\partial x}dx = xG(x)i(x,t)|_{-\infty}^{\infty} - \int_{-\infty}^{\infty} G(x)i(x,t)dx.$$

Since $xG(x)i(x,t)$ tends to 0 as $x$ approaches to $\pm\infty$, $\int_{-\infty}^{\infty} x\frac{\partial G(x)i(x,t)}{\partial x}dx = -\int_{-\infty}^{\infty} G(x)i(x,t)dx$. Using the property of the the integration of $e^x$ under a normal distribution of $x$, we evaluate the integral

$$\int_{-\infty}^{\infty} G(x)i(x,t)dx = I(t)\int_{-\infty}^{\infty} G(x)f(x;\mu_t,\sigma_F)dx = I(t)G(\mu_t + \frac{\sigma_F^2}{2}).$$

And pathogen load from initial infection is obtained by

$$\int_{-\infty}^{\infty} x\beta(t)S(t)f_0(x;\mu_0,\sigma_0)dx = \mu_0\beta_0 Z(t)S(t).$$

Then, the final form of the dynamics of total pathogen load is

$$\frac{dP(t)}{dt} = \mu_0\beta_0 Z(t)S(t) + I(t)G(\mu_t + \frac{\sigma_F^2}{2}) - I(t)[\mu_t(l + d(\alpha_t)) + \sigma_F^2 ab^{\alpha_t} lnb],$$

where $\alpha_t = \mu_t + \frac{lnb\sigma_F^2}{2}$.

The expression of environmental free-living zoospores is given by:

$$\begin{aligned}\frac{dZ}{dt} &= \underbrace{-Z(t)d_z}_{\text{Zoospore loss}} + \underbrace{\lambda\int_{-\infty}^{\infty} e^x I(x,t)\,dx}_{\text{Shedding}} \\ &= -Z(t)d_z + \lambda I(t)e^{\mu_t + \frac{\sigma_F^2}{2}}.\end{aligned}$$

The final form for the reduced dimension tracking the first moment of PDE model is presented as below:

$$\begin{aligned}\frac{dS(t)}{dt} &= Recruit(H(t)) - d_0 S(t) - \beta_0 Z(t)S(t) + lI(t),\\ \frac{dI(t)}{dt} &= \beta_0 Z(t)S(t) - lI(t) - d(\alpha_t)I(t),\\ \frac{dP(t)}{dt} &= \mu_0\beta_0 Z(t)S(t) + I(t)G\left(\mu_t + \frac{\sigma_F^2}{2}\right) - I(t)[\mu_t(l + d(\alpha_t)) + \sigma_F^2 ab^{\alpha_t} lnb],\\ \frac{dZ(t)}{dt} &= -Z(t)d_z + \lambda I(t)e^{\mu_t + \frac{\sigma_F^2}{2}},\end{aligned}$$

where $\mu_t = \frac{P}{I}$ and $\alpha_t = \mu_t + \frac{lnb\sigma_F^2}{2}$.

##### Section S1.2.2 Equilibrium

For our fixed variance model, the equilibrium mean load $x^* = \mu_t$ is achieved at time $t$ when

$$\begin{aligned} 0 &= \mu_0(l + d(\alpha_t)) + G(\theta_t) - \mu_t(d(\alpha_t) + l) - \sigma_F^2 ab^{\alpha_t} lnb \\ &= \mu_0(l + d(\alpha_t)) + G(\theta_t) - \int_{-\infty}^{\infty} x(l + d(x))i(x,t)dx. \end{aligned}$$

Where $\mu_t = \frac{P}{I} = \mu^*$, $\theta_t = \mu_t + \frac{\sigma_F^2}{2}$, and $\alpha_t = \mu_t + \frac{lnb\sigma_F^2}{2}$.

The equilibrium mean load $\mu^*$ can be interpreted as a weighted average between the load due to pathogen growth on a host and the load due to initial infection, where the relative contributions depend on survival and loss of infection. When loss of infection and mortality is high, mean pathogen load is again equal to the initial infection load $\mu_0$. However, as loss of infection and mortality becomes increasingly rare, the mean load at equilibria is approaching to pathogen load carrying capacity.

##### Section S1.2.3 Derivations of $R_0$

We follow the next-generation matrix method to derive the basic reproduction number $R_0$ (*1*, *2*). This approach constructs two matrices: **F**, which represents the rate of new infections, and **V**, which represents transitions among infected compartments. Both matrices are composed of partial derivatives evaluated at the disease-free equilibrium $S^*$. The derivations of $R_0$ for the second order approximation ODE model follows the same procedure.

The **F** matrix contains the partial derivatives of the new infection terms with respect to the infected compartments:

$$\mathbf{F} = \begin{pmatrix} 0 & 0 & \beta_0 S^* \\ 0 & 0 & \mu_0\beta_0 S^* \\ 0 & 0 & 0 \end{pmatrix}.$$

The **V** matrix contains the partial derivatives of the transfer terms between infected compartments:

$$\mathbf{V} = \begin{pmatrix} \frac{\partial V_I}{\partial I(t)} & \frac{\partial V_I}{\partial P(t)} & \frac{\partial V_I}{\partial Z(t)} \\ \frac{\partial V_P}{\partial I(t)} & \frac{\partial V_P}{\partial P(t)} & \frac{\partial V_P}{\partial Z(t)} \\ \frac{\partial V_Z}{\partial I(t)} & \frac{\partial V_Z}{\partial P(t)} & \frac{\partial V_Z}{\partial Z(t)} \end{pmatrix}$$

Specifically,

$$\begin{aligned}
V_I &= lI(t) + d(\alpha_t)I(t), \\
V_P &= -I(t)G\left(\mu_t + \frac{\sigma_F^2}{2}\right) + I(t)[\mu_t(l + d(\alpha_t)) + \sigma_F^2 ab^{\alpha_t} lnb], \\
V_Z &= Z(t)d_z - \lambda I(t)e^{\mu_t + \frac{\sigma_F^2}{2}}.
\end{aligned}$$

Once the $\mathbf{F}$ and $\mathbf{V}$ matrices are specified, the basic reproduction number $R_0$ is defined as the dominant eigenvalue (spectral radius) of the matrix product $\mathbf{FV}^{-1}$ evaluated at the disease free equilibrium.

#### Section S1.3 2nd order approximation

We derive a second-order approximation of our PDE model, which tracks both the mean and variance of pathogen load under the assumption of a lognormal distribution. We track five state variables $S(t)$ - population of susceptible individuals, and $I(t)$ - population of infected population, $P(t)$ - total pathogen load in the infected population (first moment), $Q(t)$ - second moment of pathogen load, and $Z(t)$ - free-living zoospores in the environments. The second moment of pathogen load is defined as: $Q(t) = \int_{-\infty}^{\infty} x^2 I(x,t)dx$. The variance of the lognormally distributed pathogen load, $\sigma_t^2$, is given by: $\sigma_t^2 = \mathbb{E}[x^2] - (\mathbb{E}[x]^2) = \frac{Q(t)}{I(t)} - \mu_t^2$. Changes of the second moment of pathogen load is given by:

$$\begin{aligned}
\frac{dQ(t)}{dt} &= \frac{d}{dt}\int_{-\infty}^{\infty} x^2 i(x,t)dx \\
&= \int_{-\infty}^{\infty} x^2 \frac{\partial i(x,t)}{\partial t} dx \\
&= \underbrace{\int_{-\infty}^{\infty} x^2 \beta_0 Z(t)S(t)f_0(x;\mu_0,\sigma_0)dx}_{\text{Transmission}} - \underbrace{\int_{-\infty}^{\infty} x^2 \frac{\partial G(x)i(x,t)}{\partial x}dx}_{\text{Within-host pathogen growth}} - \underbrace{\int_{-\infty}^{\infty} x^2(l + d(x))i(x,t)dx}_{\text{Recovery and mortality}}.
\end{aligned}$$

Integrating the first term, we obtain:

$$\int_{-\infty}^{\infty} x^2 \beta_0 Z(t)S(t)f_0(x;\mu_0,\sigma_0)dx = (\sigma_0^2 + \mu_0^2)\beta_0 Z(t)S(t).$$

For the second term, we use integration by parts:

$$\int_{-\infty}^{\infty} x^2 \frac{\partial G(x)i(x,t)}{\partial x}dx = x^2 G(x)i(x,t)|_{-\infty}^{\infty} - \int_{-\infty}^{\infty} 2xG(x)i(x,t)dx.$$

$x^2G(x)i(x,t)$ tends to 0 as $x$ approaches to $\pm\infty$ as log pathogen load is normally distributed and decays much more rapidly than $x^2G(x)$. We have

$$\int_{-\infty}^{\infty} x^2 \frac{\partial G(x)i(x,t)}{\partial x} dx = -\int_{-\infty}^{\infty} 2xG(x)i(x,t)dx$$
$$= -2I(t)\int_{-\infty}^{\infty} xG(x)f(x;\mu_t,\sigma_t)dx$$
$$= -2I(t)[(\mu+\sigma_t^2)G(\mu+\frac{\sigma_t^2}{2}) - \sigma_t^2 c].$$

The integration of the third term is given by

$$\int_{-\infty}^{\infty} x^2(l+d(x))i(x,t)dx = \int_{-\infty}^{\infty} x^2 li(x,t)dx + \int_{-\infty}^{\infty} x^2 d(x)i(x,t)dx$$
$$= (\sigma_t^2+\mu_t^2)lI(t) + \int_{-\infty}^{\infty} x^2 d(x)i(x,t)dx.$$

Again, we use the moment-generating function to solve the integration:

$$\int_{-\infty}^{\infty} x^2 d(x)i(x,t)dx = I(t)\int_{-\infty}^{\infty} x^2(d_0+ab^x)i(x,t)dx$$
$$= I(t)[(\sigma_t^2+\mu_t^2)d_0 + a\mathbb{E}[x^2e^{xlnb}]].$$

To derive $\mathbb{E}(x^2e^{xlnb})$, we take the second derivative of $\mathbb{E}(e^{xlnb})$, where $lnb = m$. Then, we have $\mathbb{E}(e^{xlnb}) = \mathbb{E}(e^{mx})$, and

$$\mathbb{E}(x^2e^{mx}) = \frac{d^2\mathbb{E}(e^{mx})}{d^2m}.$$

Since $x \sim \mathcal{N}(\mu_t,\sigma_t)$, we know that $\mathbb{E}(e^{mx}) = e^{m\mu_t+\frac{m^2\sigma_t^2}{2}}$, and its second derivative is:

$$\mathbb{E}(x^2e^{mx}) = \frac{d^2\mathbb{E}(e^{mx})}{d^2m}$$
$$= \mathbb{E}(e^{mx})\sigma_t^2 + \mathbb{E}(e^{mx})(\mu_t+m\sigma_t^2)^2$$
$$= (\sigma_t^2+(\mu_t+m\sigma_t^2)^2)e^{m\mu_t+\frac{m^2\sigma_t^2}{2}}.$$

Evaluate at $m = lnb$, we have

$$\mathbb{E}(x^2e^{xlnb}) = (\sigma_t^2+(\mu_t+lnb\sigma_t^2)^2)e^{lnb\mu_t+\frac{lnb^2\sigma_t^2}{2}}.$$

Substituting back, we obtain:

$$\int_{-\infty}^{\infty} x^2 d(x)I(x,t)dx = I(t)\int_{-\infty}^{\infty} x^2(d_0+ab^x)i(x,t)dx$$
$$= I(t)[(\sigma_t^2+\mu_t^2)d_0 + a(\sigma_t^2+(\mu_t+lnb\sigma_t^2)^2)e^{lnb\mu_t+\frac{lnb^2\sigma_t^2}{2}}]$$
$$= I(t)[(\sigma_t^2+\mu_t^2)d(\alpha_t) + 2lnb\sigma_t^2\alpha_t ab^{\alpha_t}],$$

where we define $\mu_t = \frac{P(t)}{I(t)}$ and $\alpha_t = \mu_t + \frac{lnb\sigma_F^2}{2}$.

The expression for the time derivative of the second moment $Q(t)$ of pathogen load is:

$$\frac{dQ(t)}{dt} = (\sigma_0^2 + \mu_0^2)\beta_0 Z(t)S(t) + 2I(t)[(\mu_t + \sigma_t^2)G(\mu_t + \frac{\sigma_t^2}{2}) - \sigma_t^2 c] \\ - (\sigma_t^2 + \mu_t^2)I(t)(l + d(\alpha_t)) - 2lnb\sigma_t^2\alpha_t ab^{\alpha_t}I(t).$$

The final form of the second order approximation is:

$$\frac{dS(t)}{dt} = \text{Recruit}(H(t)) - d_0 S(t) - \beta_0 Z(t)S(t) + lI(t),$$
$$\frac{dI(t)}{dt} = \beta_0 Z(t)S(t) - lI(t) - d(\alpha_t)I(t),$$
$$\frac{dP(t)}{dt} = \mu_0\beta_0 Z(t)S(t) + I(t)G\left(\mu_t + \frac{\sigma_t^2}{2}\right) - I(t)\left[\mu_t(l + d(\alpha_t)) + \sigma_t^2 ab^{\alpha_t}\ln b\right],$$
$$\frac{dQ(t)}{dt} = (\sigma_0^2 + \mu_0^2)\beta_0 Z(t)S(t) + 2I(t)\left[(\mu_t + \sigma_t^2)G\left(\mu_t + \frac{\sigma_t^2}{2}\right) - \sigma_t^2 c\right] \\ - (\sigma_t^2 + \mu_t^2)I(t)(l + d(\alpha_t)) - 2\ln(b)\sigma_t^2\alpha_t ab^{\alpha_t}I(t),$$
$$\frac{dZ(t)}{dt} = -Z(t)d_z + \lambda I(t)e^{\mu_t + \frac{\sigma_t^2}{2}}.$$

The mean of pathogen load is given by $\mu_t = \frac{P(t)}{I(t)}$ and the variance is $\sigma_t^2 = \frac{Q(t)}{I(t)} - \mu_t^2$.

##### Section S1.3.1 Equilibrium

The equilibrium mean load $x^* = \mu_t$ and variance $y^* = \sigma_t^2$ is achieved at time $t$ when:

$$0 = \mu_0(l + d(\alpha_t)) + G(\theta_t) - \mu_t(d(\alpha_t) + l) - \sigma_t^2 ab^{\alpha_t}lnb \\ = \mu_0(l + d(\alpha_t)) + G(\theta_t) - \int_{-\infty}^{\infty} x(l + d(x))i(x,t)dx,$$

and the equilibrium condition for the second moment is:

$$0 = (\mu_0^2 + \sigma_0^2)(l + d(\alpha_t)) + 2[(\mu_t + \sigma_t^2)G(\theta_t) - \sigma_t^2 r_{max}] - (\sigma_t^2 + \mu_t^2)(l + d(\alpha_t)) - 2lnb\sigma_t^2\alpha_t ab^{\alpha} \\ = (\mu_0^2 + \sigma_0^2)(l + d(\alpha_t)) - \int_{-\infty}^{\infty} x^2 \frac{\partial G(x)i(x,t)}{\partial x}dx - \int_{-\infty}^{\infty} x^2(l + d(x))i(x,t)dx.$$

Here, the terms are defined as: $\mu_t = \frac{P}{I} = \mu^*$, $\theta_t = \mu_t + \frac{\sigma_t^2}{2}$, and $\alpha_t = \mu_t + \frac{lnb\sigma_t^2}{2}$.

### Section S2 Parameters

Table S1: Parameter values, units, and sources. Parameters are based on the California red-legged frog (*Rana draytonii*) when available, and obtained from the mountain yellow-legged frog (*Rana muscosa*) when noted.

| Parameters | Value | Details | Reference |
|---|---|---|---|
| Background mortality rate (frogs/day) | $d_0 = 0.001$ | Corresponding to a annual adult survivorship of 0.686 | Licht (3) |
| Recruitment (frogs/day) and density dependence | $r = 0.013$, $\gamma = 0.033$ | Each adult produces 1,500 eggs annually, with survivorship rates of 0.025 in the first year, 0.25 in the second year, and 0.52 in the third year. | Licht (3) and Doubledee *et al.* (4) |
| Load-dependent mortality (frogs/day) | $a = 0.000025$, $b = e^{1.1}$ | Fifty percent survivorship occurs at a log pathogen load of 7.6. | Bradley *et al.* (5) |
| Within-host pathogen growth (zoospores/day) and carrying capacity (zoospores) | $r_{max} = 0.664$, $K = e^{12}$ | Approximate the density-dependent growth curve at 20 $°C$ (R. muscosa). | Wilber *et al.* (6) |
| Recovery rate (frogs/day) | $l = 0.0032$ | Assuming log pathogen load 6 at 20 $°C$ (R. muscosa). | Wilber *et al.* (6) |
| Zoospore loss rate (zoospores/day) | $d_z = 4.658$ | Total encystment and mortality rate at 23 $°C$ | Woodhams *et al.* (7) |
| Shedding rate (zoospores/day) | $\lambda = 0.052$ | | |
| Transmission coefficiency (frogs/zoospore/day) | $\beta = 1.8 \times 10^{-4}$ | (R. muscosa) | Wilber *et al.* (8) |
| Initial pathogen load (ln(zoospores)) | $\mu_0 = 3.382$, $\sigma_0 = 2.708$ | At 20 $°C$ (R. muscosa) | Wilber *et al.* (6) |

#### Section S3 Results

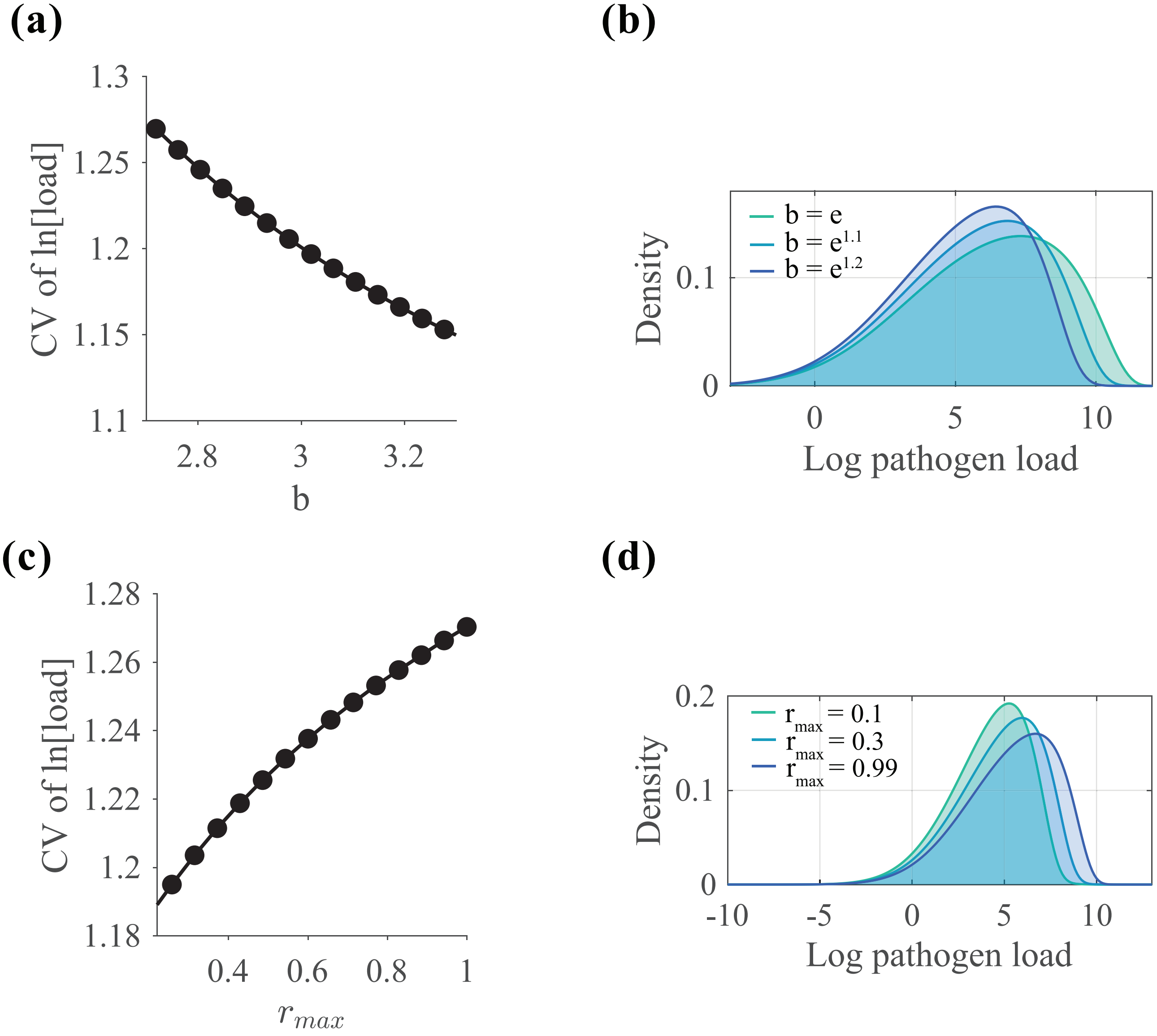


Figure S2: Infection processes influencing pathogen aggregation based on full PDE model with deterministic within-host pathogen growth. (a) the steepness of load-dependent mortality (reflecting host tolerance) affects the aggregation of pathogen load, measured by the coefficient of variation [CV]; (b) pathogen load distributions under varying levels of load-dependent mortality; (c) the rate of within-host pathogen growth influences aggregation; (d) pathogen load distributions under different within-host growth rates.

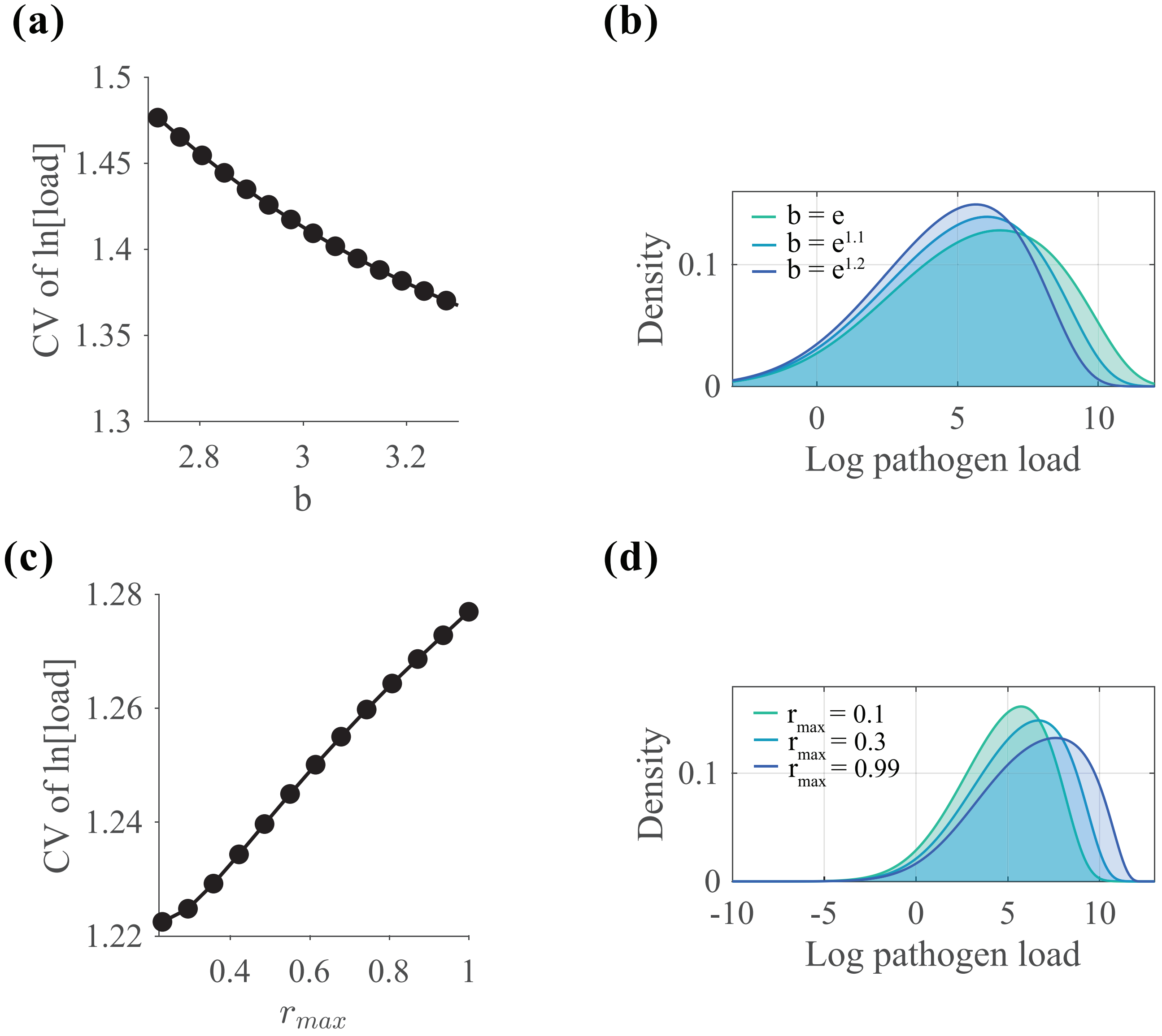


Figure S3: Infection processes influencing pathogen aggregation based on full PDE model with stochastic within-host pathogen growth. (a) the steepness of load-dependent mortality (reflecting host tolerance) affects the aggregation of pathogen load, measured by the coefficient of variation [CV]; (b) pathogen load distributions under varying levels of load-dependent mortality; (c) the rate of within-host pathogen growth influences aggregation; (d) pathogen load distributions under different within-host growth rates.

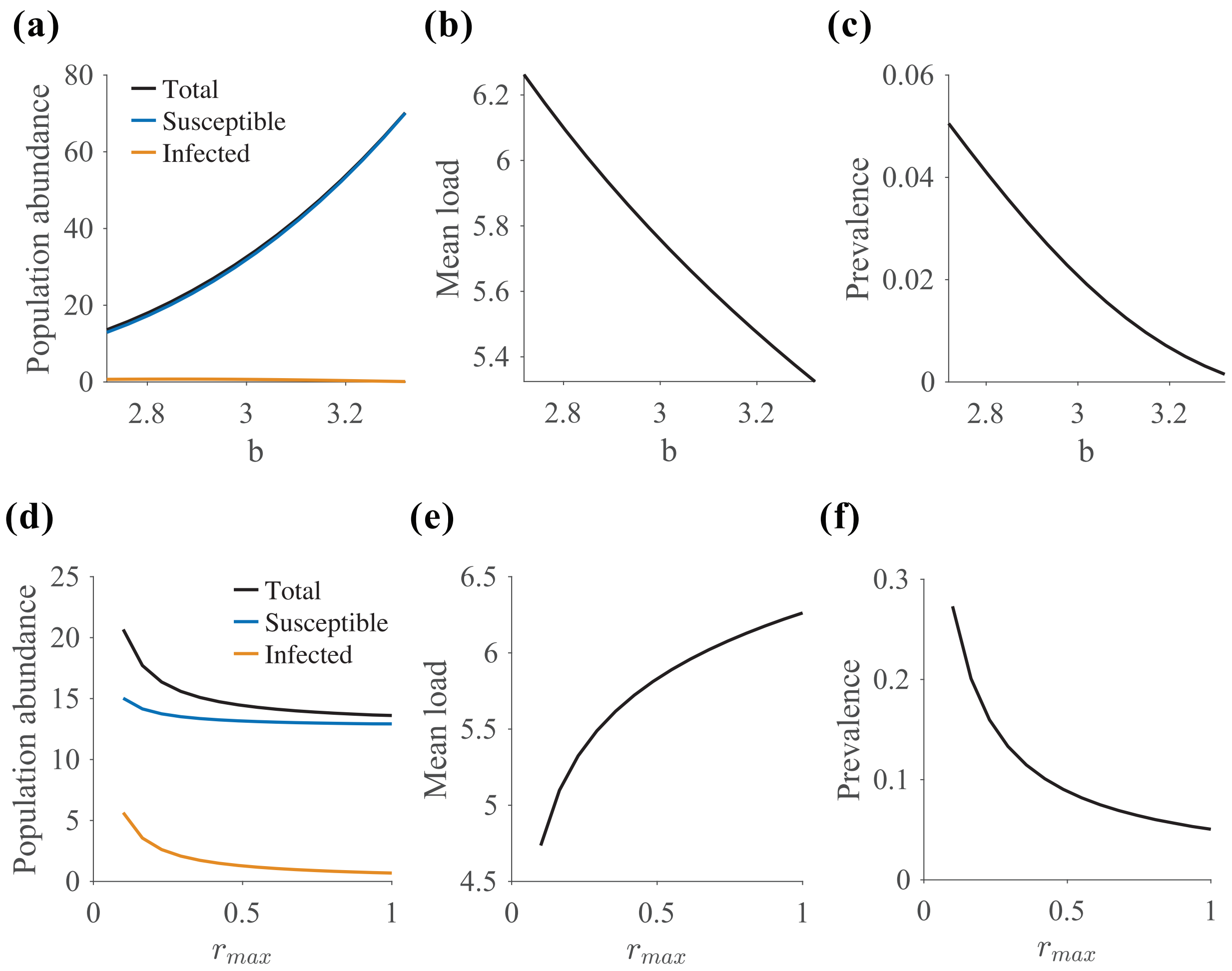


Figure S4: Effects of infection processes on host population dynamics based on PDE model with deterministic within-host pathogen growth. Panels show the effects of (a–c) load-dependent mortality, and (d–f) within-host pathogen growth on host population suppression, mean pathogen load of infected individuals, and disease prevalence.

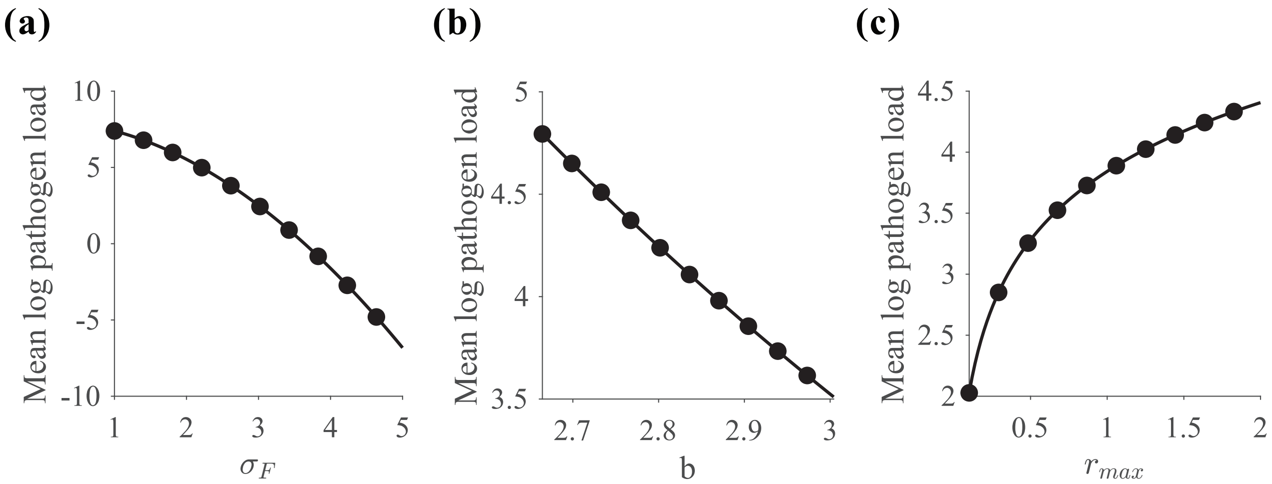


Figure S5: Effects of (a) pathogen aggregation, (b) steepness of the load-dependent mortality curve, and (c) within-host pathogen growth rate on the mean pathogen load of infected individuals, based on Model 1.

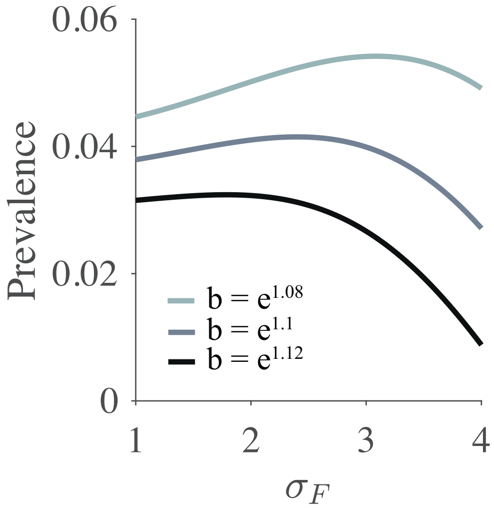


Figure S6: Effects of pathogen aggregation on disease prevalence under varying steepness levels of the load-dependent mortality curve, based on Model 1.

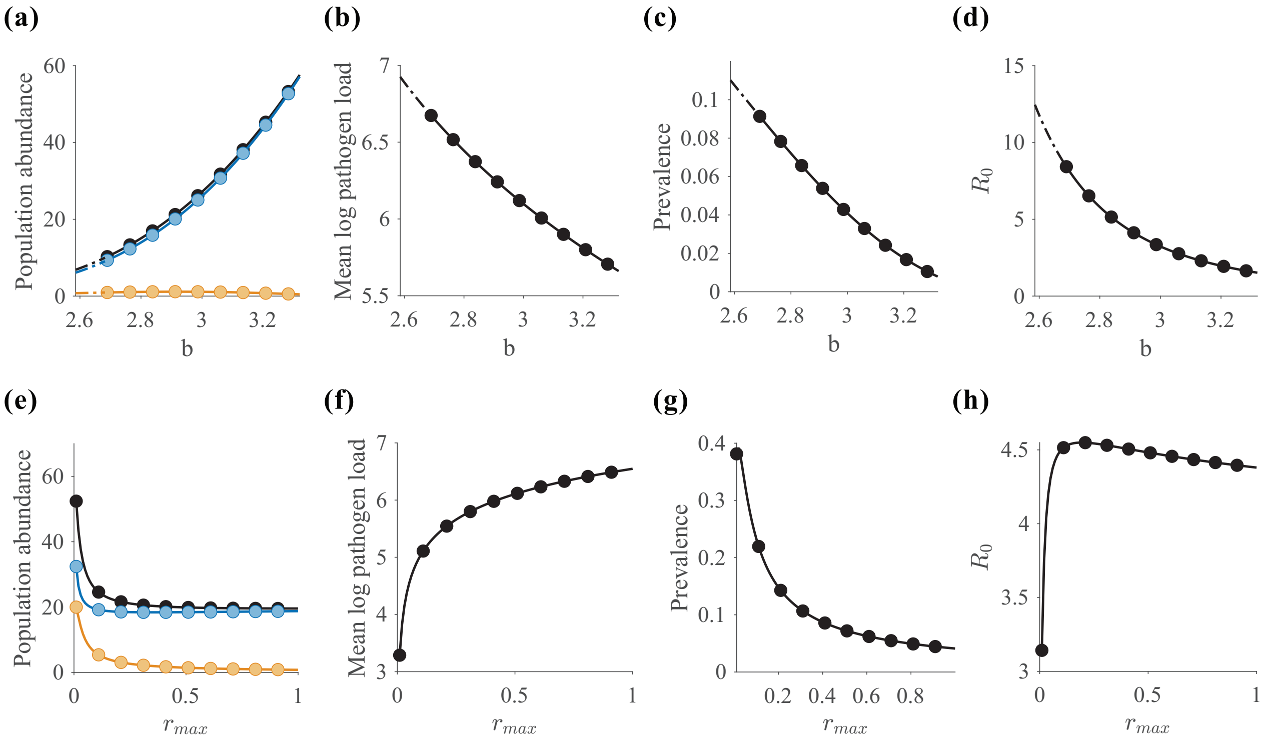


Figure S7: Effects of infection processes on host population dynamics and disease transmission potential ($R_0$) based on Model 2. Panels show the effects of (a–d) load-dependent mortality and (e–h) within-host pathogen growth on host population suppression, mean pathogen load, disease prevalence, and the basic reproduction number ($R_0$).

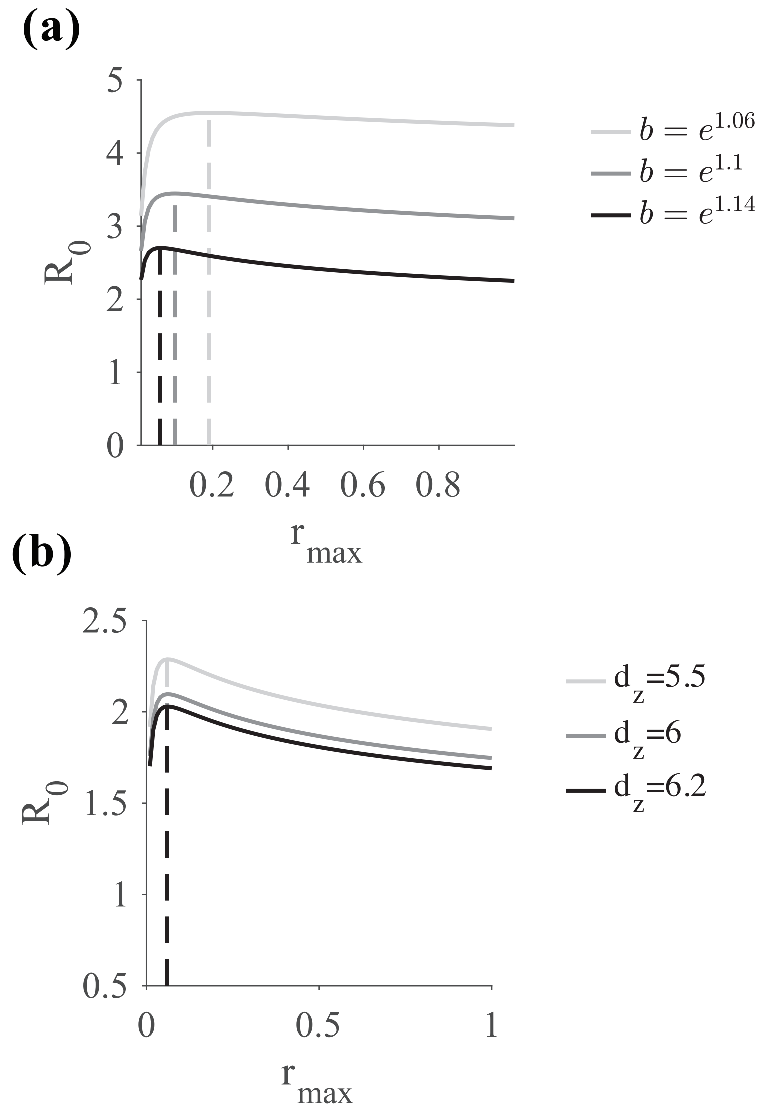


Figure S8: Factors influencing virulence–transmission trade-offs. Load-dependent mortality (a) and the loss rate of environmental zoospores (b) shape the relationship between virulence and transmission. The dashed line indicates the level of within-host pathogen growth that maximizes the basic reproduction number ($R_0$).
